# Nkx2.1 regulates the proliferation and cell fate of telencephalic astrocytes during embryonic development

**DOI:** 10.1101/065169

**Authors:** Shilpi Minocha, Delphine Valloton, Yvan Arsenijevic, Jean-René Cardinaux, Raffaella Dreier Guidi, Jean-Pierre Hornung, Cécile Lebrand

## Abstract

The homeodomain transcription factor Nkx2.1 controls cell differentiation of telencephalic GABAergic interneurons and oligodendrocytes. Here, we show that Nkx2.1 additionally regulates astrogliogenesis of the telencephalon from embryonic day (E) 14.5 to E16.5. Our work aims to identify the different mechanisms by which Nkx2.1 controls telencephalic astrogliogenesis. In *Nkx2.1^-/-^*, a drastic loss of astrocytes is observed which is not related to cell death. *In vivo* analysis using BrdU incorporation reveals that Nkx2.1 affects the proliferation of ventral neural stem cells that generate early astrocytes. *In vitro* neurosphere assays show that Nkx2.1 additionally affects the differentiation step of Nkx2.1-derived astrocytes. Chromatin immunoprecipitation and *in vitro* co-transfection studies of a Nkx2.1-expressing plasmid indicate that Nkx2.1 binds to the promoter of astroglial differentiation gene GFAP, and regulates its expression. Hence, Nkx2.1 controls astroglial production spatiotemporally in embryos by regulating stem cell division and specification of the contributing Nkx2.1^+^ precursors.

## Introduction

Proper forebrain development is carried out by coordinated and regulated developmental events involving controlled cell proliferation, differentiation, and guided migration of neuronal and glial cells. Several spatiotemporally orchestrated molecular mechanisms underlie the successful patterning of the telencephalon (Guillemot et al., 2006, Long et al., 2009, Marin and Rubenstein, 2001, Puelles et al., 2000, Schuurmans et al., 2004, Schuurmans and Guillemot, 2002, Yun et al., 2001). Both dorsal and ventral telencephalons are demarcated by specific gene expression that regulates the generation of defined neuronal and glial populations. Dorsal progenitors express homeobox genes of the empty spiracles (*Emx1*/*Emx2*), and paired homeobox (*Pax6*) families, and *atonal*-related genes Neurogenin (*Ngn*)*1*/*Ngn2* whereas the ventral progenitors are known to exhibit expression of homeobox genes of the *Nkx* (*Nkx2.1*) and distal-less (*Dlx1*/*Dlx2*) families, *Gsh1*/*2*, and *achaete*-*scute-* related gene *Mash1* (Campbell, 2003, Guillemot, 2007b, Morrow et al., 2001, Qian et al., 2000, Rubenstein et al., 1998).

Broadly, amongst the neuronal population, the glutamatergic projection neurons have been shown to be primarily generated by dorsal telencephalic progenitors whereas the GABAergic interneurons originate from the ventral telencephalic progenitors (Kriegstein and Noctor, 2004, Marin et al., 2000, Marin and Rubenstein, 2001, Molyneaux et al., 2007). Amongst the glial population, the embryonic oligodendrocytes are produced in waves from the ventral telencephalic progenitors (Kessaris et al., 2006, Kessaris et al., 2008). On the other hand, the exact timing of generation and origin of the embryonic astroglial population is still a topic of active investigation. Embryonic astrocytes have been shown to be either generated from bipotential radial glia or from progenitor cells in the subventricular zone (Levison and Goldman, 1993, Schmechel and Rakic, 1979, Guillemot, 2007a, Kriegstein and Alvarez-Buylla, 2009, Mori et al., 2005, Pinto and Gotz, 2007). In the dorsal telencephalon, astrocyte gliogenesis has been mostly documented to only occur after neurogenesis (after E17, in mice) when the bipotential radial glial cells of the dorsal pallium differentiate into astrocytes (Guillemot, 2007a, Mission et al., 1991, Rowitch and Kriegstein, 2010, Price and Thurlow, 1988, Cameron and Rakic, 1991, Lavdas et al., 1999, Schmechel and Rakic, 1979, Gotz and Huttner, 2005). Several indusium griseum (IG) glia, surrounding the CC, are also shown to originate from the radial glia of the dorsomedial pallium (Smith et al., 2006). The time of generation of some of the astrocytes occupying the CC midline region, however, is noted to be between E13 and postnatal day 2 (P2) with a peak at E14, much earlier than previously proposed (Shu et al., 2003). Furthermore, the postnatal astrocytes that occupy the cerebral cortex region are believed to originate from progenitor cells in the dorsolateral subventricular zone (SVZ) (Marshall et al., 2003). Recent evidence shows that a population of locally differentiated glia in the postnatal cortex instead constitute the primary source of astrocytes rather than the aforementioned SVZ progenitors (Ge et al., 2012). Since the glia play essential roles in guidance of forebrain commissures during embryonic brain development, hence, the detailed understanding of the point of origin(s) and exact timing of generation of the telencephalic astroglia is necessary.

Nkx2.1, a homeodomain transcription factor, was initially found to regulate the transcription of many thyroid-specific genes (Guazzi et al., 1990, Lazzaro et al., 1991, Sussel et al., 1999) and lung-specific genes (Boggaram, 2009, Hamdan et al., 1998). Furthermore, several cell cycle related genes such as *Notch1*,*E2f3*, *Cyclin B1*, *Cyclin B2* and *c-Met*, have been found to be bound by Nkx2.1 in developing embryonic lungs (Tagne et al., 2012). In the brain, it is known to control during embryonic development, the specification of GABAergic interneurons and oligodendrocytes that populate the ventral and dorsal telencephalic region (Anderson et al., 2001, Corbin et al., 2001, Kessaris et al., 2006, Kimura et al., 1996, Marin and Rubenstein, 2001, Sussel et al., 1999). Loss of Nkx2.1 leads to ventral-to-dorsal respecification of the pallidum, and causes loss of GABAergic interneurons and oligodendrocytes in the dorsal telencephalic region (Kessaris et al., 2006, Kessaris et al., 2008, Sussel et al., 1999). Recently, we showed that during embryonic development Nkx2.1 also regulates the generation of astrocytes that populate the ventral telencephalon and participates to axonal guidance in the anterior commissure (Minocha et al., 2015a, Minocha et al., 2015b). We find that this Nkx2.1-derived astrocyte population are generated from three ventral telencephalic precursor regions, namely the medial ganglionic eminence (MGE), the anterior entopeduncular area (AEP)/ preoptic area (POA), and the triangular septal nucleus (TS) (Minocha et al., 2015a, Minocha et al., 2015b).

In this study, we found that Nkx2.1-derived astrocytes populate the corpus callosum (CC) and its surrounding regions in the embryonic dorsal telencephalon. The Nkx2.1-derived astrocytes are generated from E12.5 onwards with maximal production between E14.5-to-E16.5. Interestingly, by using *Nkx2.1^-/-^* mice, we observed that the functional ineffectiveness of the mutated Nkx2.1 (mut-Nkx2.1) leads to a drastic loss of astrocytes and polydendrocytes in the entire dorsal telencephalic region at the midline. Since the aforementioned Nkx2.1-derived cell loss is not accompanied with increased cell death, we further analyzed if cell proliferation of Nkx2.1^+^ stem cells in the three ventral precursor regions (MGE, AEP/POA and TS) was affected. *In vivo* BrdU incorporation and *in vitro* neurosphere differentiation assays showed that Nkx2.1 interestingly exerts its control over astroglial generation by controlling both the proliferation and differentiation capacity of the Nkx2.1^+^ precursors. In addition, chromatin immunoprecipitation analysis indicated that the transcription factor Nkx2.1 binds to the promoter of astroglial differentiation gene, glial fibrillary acidic protein (*GFAP*). Co-transfection studies with GFAP promoter construct and tagged Nkx2.1 over-expression in HEK293 cells confirmed that Nkx2.1 could indeed regulate the expression of *GFAP* gene. Hence, Nkx2.1 regulates astroglial generation by regulating the proliferation and differentiation of the contributing Nkx2.1^+^ precursors in ventral telencephalic zones. Thus, Nkx2.1 exhibits a multilevel control over the generation and differentiation of the telencephalic astroglia by spatially coordinating the astroglial generation from the three aforementioned precursor regions and temporally restricting maximal generation between E14.5-to-E16.5. Further analysis into the complete repertoire of genes regulated by Nkx2.1 can shed light about the glial and neuronal populations that play a defining role in shaping the brain.

## Results

### Nkx2.1-derived astrocytes populate the dorsal telencephalon during development

Nkx2.1-positive (Nkx2.1^+^) progenitors of the MGE, the AEP/POA and the septal nucleus contribute towards the production of embryonic GABAergic interneurons and oligodendrocytes that populate the ventral and dorsal telencephalon (Anderson et al., 2001, Corbin et al., 2001, Kessaris et al., 2006, Kimura et al., 1996, Marin and Rubenstein, 2001, Sussel et al., 1999). Our recent results have shown that Nkx2.1 additionally regulates the production of astrocytes and polydendrocytes that populate the ventral telencephalon (Minocha et al., 2015a, Minocha et al., 2015b). Here, interestingly, further immunostaining against the subpallial transcription factor Nkx2.1 revealed strong expression in several differentiated cells within and surrounding the CC in the dorsal telencephalon from E16.5 to E18.5 too (n=8; Fig. 1a-g). To further differentiate the Nkx2.1^+^ cell types in the CC region, we made use of several cell-type specific transgenic strains and immunohistochemical analysis for neuronal and glial makers.

**Figure 1.**
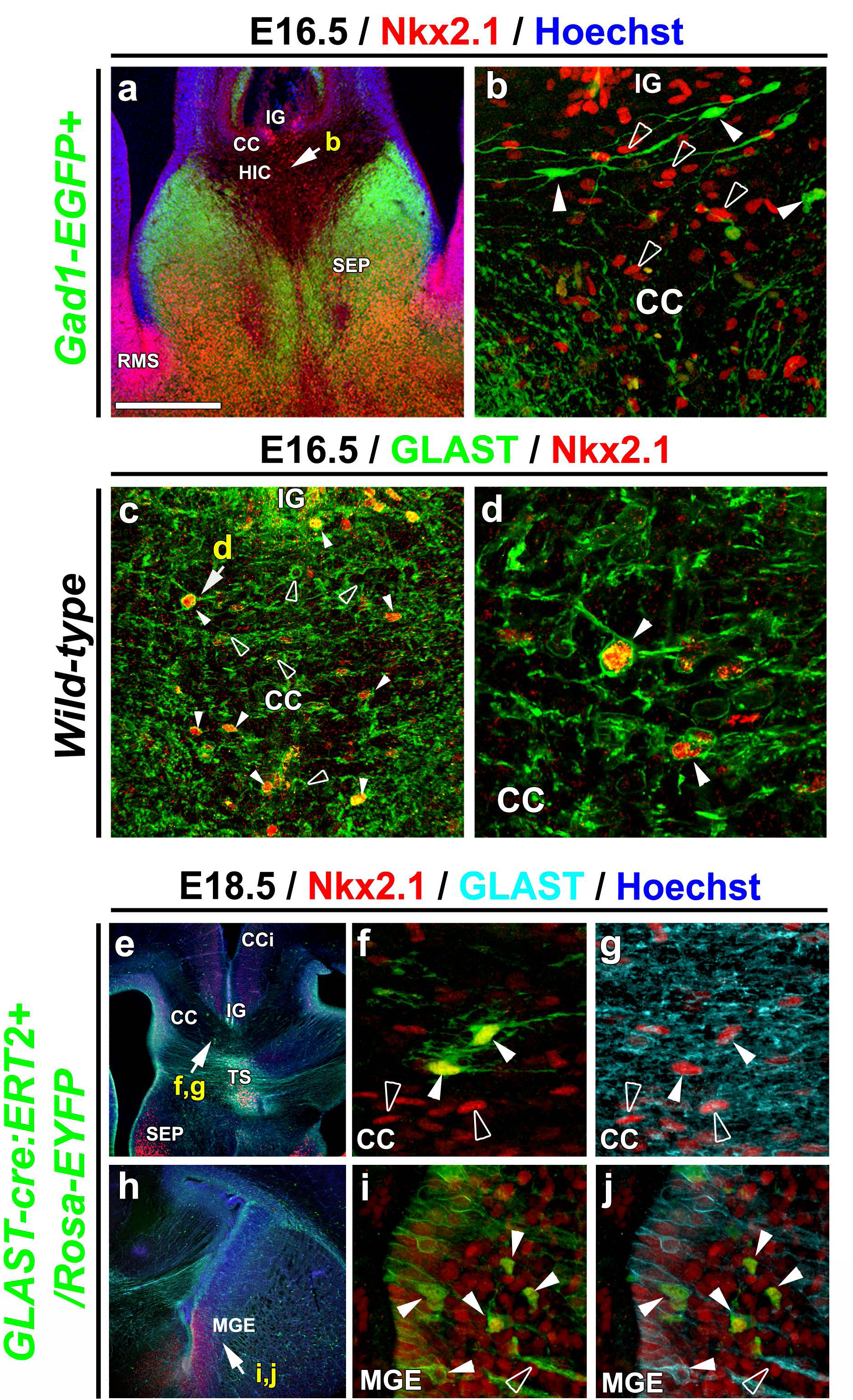
The Nkx2.1-positive cells of the CC are glial cells. **(a-d)** Double immunohistochemistry for the GFP and Nkx2.1 **(a-b)** on coronal CC sections from *Gad1-EGFP^+^* mice at E16.5 (n=4), and for GLAST and Nkx2.1 (n=2) **(c-d)** on coronal CC sections from wild-type mice at E16.5. **(e-j)** Triple immunohistochemistry for the EYFP, Nkx2.1 and GLAST on coronal CC **(e-g)** and MGE ( **h-j)** sections from *GLAST-Cre:ERT2^+^/Rosa-EYFP* mice at E18.5 (n=5). Cell nuclei were counterstained in blue with Hoechst **(a, e** and **h)**. Colocalization between the green and the red channel is highlighted in yellow **(b, c, d, f, i** and **j)**. **b, d, f, g, i,** and **j** are higher power views of the CC and MGE region indicated by an arrow in **a, c, e** and **h**, respectively. **(a-d)** At E16.5, several Nkx2.1^+^ (red) nuclei were observed in the medial part of the CC (open arrowheads in **b**). Most of the *Gad1-EGFP*^+^interneurons (green) populating this region were not labelled by Nkx2.1 (solid arrowhead in **b)**. At this age, however, colocalization revealed that most of the Nkx2.1-expressing nuclei co-expressed astroglial markers like GLAST (solid arrowheads in **c** and **d)**.**(e-j)** The Cre-mediated recombination was initiated under the control of the tamoxifen-inducible GLAST promoter at E14, and the GLAST-derived astroglia were visualized (in green) with the EYFP signal. Some GLAST^+^ astroglial cells (in light blue) of the CC **(e-g)** and the MGE **(i-j)** co-expressed Nkx2.1 (in red, solid arrowheads). Some of the GLAST^+^/Nkx2.1^+^ glia were not labelled by the EYFP signal and might have been generated before the recombination was induced (open arrowheads in **f** and **g)**. **(CC)** corpus callosum; **(CCi)** cingulate cortex; **(IG)** induseum griseum; **(HIC)** hippocampal commissure; **(MGE)** medial ganglionic eminence; **(RMS)** rostral migratory stream; **(SEP)** septum; **(TS)** triangular septal nucleus. Bar = 675 µm in **e** and **h**; 450 µm in **a**; 67 µm in **b** and **c**; 40 µm in **f**, **g**, **i** and **j**; 30 µm in **d**.

Firstly, to identify the GABAergic interneurons, we made use of the *Gad1-EGFP* knock-in mice, which express the green fluorescent protein (EGFP) in GAD67^+^ GABAergic interneurons (Tamamaki et al., 2003) in combination with immunostaining against Nkx2.1. Coherent with previous observations showing down-regulation of Nkx2.1 expression in dorsal telencephalic GABAergic interneuron population (Nobrega-Pereira et al., 2008), at E16.5, none of the *Gad1-GFP*^+^ interneurons of the CC and dorsal surrounding areas expressed Nkx2.1 (n=4; Fig. 1a-b; solid arrowheads in 1b).

Secondly, to ascertain if Nkx2.1^+^ cells corresponded to polydendrocytes, we made use of the *Cspg4-Cre^+^/Rosa-EYFP* mice that express the yellow fluorescent protein (EYFP) in NG2^+^ polydendrocytes (Nishiyama et al., 2009, Nishiyama et al., 2002, Minocha et al., 2015a). We found that the EYFP was expressed by Nkx2.1^+^ progenitors of the MGE (n=3; Figure 1-figure supplement 1a-c, solid arrowheads in b-c). However, the EYFP^+^ polydendrocytes stopped to express the Nkx2.1 protein outside ventral germinal zones and within the dorsal telencephalon (n=3; Figure 1-figure supplement 1g-i, open arrowheads in h-i). Hence, to further delineate the profile of Nkx2.1^+^ cells in the CC and the surrounding areas, we performed immunostaining in wild-type embryos, against Nestin and GLutamate and ASpartate Transporter (GLAST) that are specific for post-mitotic astrocytes within the embryonic CC white matter (Shu et al., 2003). Interestingly, from E16.5 to E18.5, 2/3^rd^ of Nkx2.1^+^ cells of the CC and surrounding regions were found to be GLAST^+^ astrocytes (n=11; Fig. 1c-d, 1f-g, 2a-b, 2e, Figure 1-figure supplement 1h-i, solid arrowheads). After postnatal day 0 (P0), however, Nkx2.1 was strongly down-regulated and not detected anymore by immunohistochemistry in the astrocytes of the dorsal telencephalon (n=8, not shown). Nkx2.1 expression was never detected in any radial glial precursor cells of the glial wedge (GW) and of the dorsal telencephalic ventricular zone labeled for the aforementioned astrocytic markers (n=11; Figure 1-figure supplement 2).

**Figure 2.**
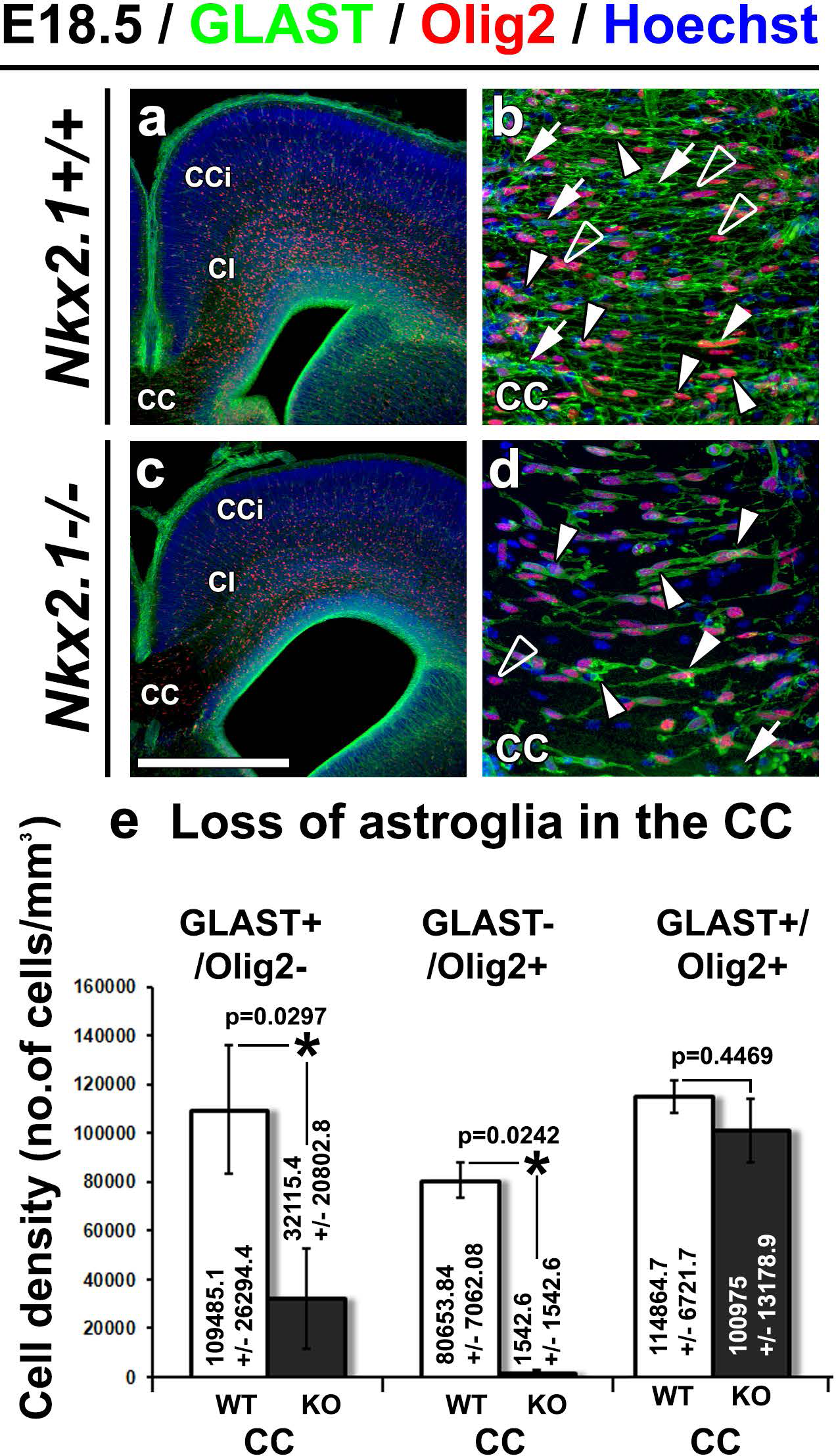
Loss of GLAST^+^/Olig2^-^ astroglia and GLAST-/Olig2^+^ polydendrocytes in the CC of *Nkx2.1^-/-^* mutant mice. **(a-d)** Double immunohistochemical staining for Olig2 and GLAST on CC coronal sections from wild-type *Nkx2.1^+/+^* (n=4) **(a-b)** and *Nkx2.1^-/-^* (n=2) **(c-d)** mice at E18.5. Cell nuclei were counterstained in blue with Hoechst **(a-d)**. **b** and **d** are higher power views of the CC seen in **a** and **c**, respectively. **(b** and **d)** In the CC midline of *Nkx2.1^-/-^* mice, there was a severe loss of GLAST^+^/Olig2^-^ astroglia (arrows) and GLAST^-^/Olig2^+^ polydendrocytes (open arrowheads) but not of GLAST^+^/Olig2^+^ astroglia (solid arrowheads) compared to the wild-type mice. **(e)** Bars (mean ± SEM from a sample of n=3 wild-type and n=3 *Nkx2.1^-/-^*) represent the density (number of cells/mm^3^) of GLAST^+^/Olig2^-^, GLAST^-^/Olig2^+^ and GLAST^+^/Olig2^+^ glial cells in the CC of *Nkx2.1^-/-^* (KO) compared to wild-type (WT) mice at E18.5. The quantification confirms the significant decrease of the GLAST^+^/Olig2^-^ astroglia (p-value=0.0297) and GLAST^-^/Olig2^+^polydendrocytes (p-value=0.0242) and no change in GLAST^+^/Olig2^+^ astroglia (p-value=0.4469) in the *Nkx2.1^-/-^* CC compared to WT mice. **(CC)** corpus callosum; **(CCi)** cingulate cortex; **(CI)** cingulate bundle. Bar = 675 µm in **a** and **c**; 100 µm in **b** and **d**.

Further analyses using tamoxifen-inducible *GLAST-Cre ERT^TM^*/*Rosa26-EYFP* mice, displayed the presence of many EYFP^+^ early astrocytes outside the germinal zones and also within the CC and surrounding area from E16.5 to E18.5 (n=5; Fig. 1e-j). Many of these early astrocytes visualized by the EYFP and GLAST co-staining showed Nkx2.1 expression as well (n=5; CC in Fig. 1f-g and MGE in 1i-j, solid arrowheads).

Therefore, our previous and current extensive immunohistochemical analyses in combination with different transgenic strains reveal that the CC and the surrounding regions are populated with various Nkx2.1-derived glial cell types (summarized in Figure 1-figure supplement 3) (Minocha et al., 2015b). Presence or absence of Nkx2.1 protein expression primarily divides the Nkx2.1-derived glial classes into two major subtypes in both dorsal and ventral telencephalon — astrocyte-like or polydendrocyte-like. The Nkx2.1-derived astrocyte-like population is further sub-divided into two populations: GLAST^+^/GFAP^+^/Nkx2.1^+^ (orange) and GLAST^+^/GFAP^-^/Nkx2.1^+^ (green) (Figure 1-figure supplement 3). The polydendrocyte-like population is further sub-divided into two populations: S100β^+^/NG2^+^/Olig2^+^/Nkx2.1^-^ (red) and S100β^-^/NG2^+^/Olig2^+^/Nkx2.1^-^ (brown) populations (Figure 1-figure supplement 3). Interestingly, a subpopulation of GLAST^+^ astrocyte-like population within the telencephalon (blue) are not Nkx2.1-derived and express Olig2^+^ cells (n=4; Fig. 2a-b and 2e; Figure 1-figure supplement 3) (Minocha et al., 2015b).

Nkx2.1-derived astrocyte-like cells populate the CC region toward the end of embryonic period. Use of tamoxifen-inducible *GLAST-Cre ERT^TM^*/*Rosa26-EYFP* mice indicated the generation of several Nkx2.1^+^/GLAST^+^/EYFP^+^ astrocytes at a period beginning from E14.5 onwards as the tamoxifen injection was delivered at E14.5 (n=5; Fig. 1e-g). In addition, in order to further decipher the exact timing of generation of these Nkx2.1^+^ glial cells that occupy the CC region, we administered 5-bromo-2’-deoxyuridine (BrdU) injections to WT pregnant females bearing embryos at E12.5 (n=2), at E14.5 (n=2), and at E16.5 (n=2), and followed the extent of BrdU incorporation at E18.5 by CC astrocytes co-expressing Nkx2.1 and astroglial markers, GLAST or GFAP (Fig. 3). Combined immunostaining revealed the presence of few BrdU^+^/Nkx2.1^+^/GLAST^+^ and BrdU^+^/GFAP^+^ embryonic astrocytes in the CC when BrdU injection was delivered at E12.5 (Fig. 3a-c and j, solid arrowheads). The bulk of the BrdU^+^/Nkx2.1^+^/GLAST^+^ and BrdU^+^/GFAP^+^ embryonic astrocytes was observed when BrdU injection was delivered at E14.5 (Fig. 3d-f and k, solid arrowheads), and at E16.5 (Fig. 3h-i and l, solid arrowheads). Therefore, the majority of Nkx2.1-derived astrocytes occupying the CC region are produced between E14.5 to E16.5.

**Figure 3.**
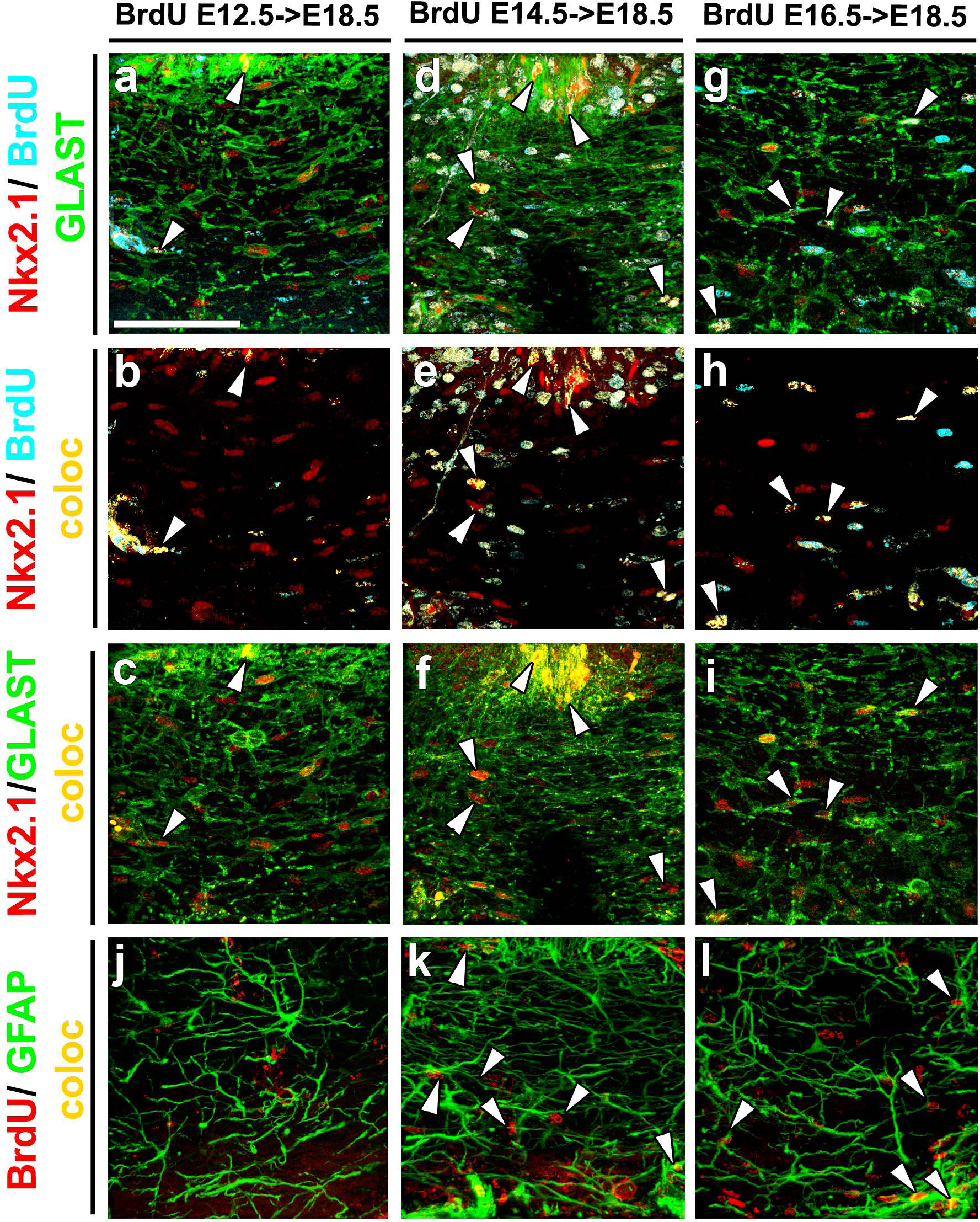
Nkx2.1-positive astroglia of the CC are generated between E14.5 and E16.5. **(a-i)** Triple immunohistochemistry for Nkx2.1, 5-bromo-2’-deoxyuridine (BrdU), and GLAST, and **(j-l)** double immunohistochemistry for BrdU and GFAP on CC coronal sections from wild-type mice brains at E18.5 injected at E12.5 (n=2) **(a-c** and **j)**, E14.5 (n=2) **(d-f** and **k)** and E16.5 (n=2) **(g-i** and **l).** **(a-i)** At E18.5, several GLAST^+^ astroglial cells (green) expressing Nkx2.1 (red) are present in the CC midline. **(b, e** and **h)** Colocalization between the blue (BrdU) and the red (Nkx2.1) channel is highlighted in yellow. **(c, f** and **i)** Colocalization between the green (GLAST) and the red (Nkx2.1) channel is highlighted in yellow. The solid arrowheads point towards the Nkx2.1^+^/GLAST^+^/BrdU^+^ cells revealing that the bulk of division for the Nkx2.1^+^ astroglial cells of the CC occurs between E14.5 **(e)** and E16.5 **(h)**. **(j-l)** Numerous GFAP^+^ astroglial cells (in green) are present in the CC midline. Colocalization between the green (GFAP) and the red (BrdU) channel is highlighted in yellow. The solid arrowheads are pointing on the GFAP^+^/BrdU^+^ cells depicting that the bulk of division for the GFAP^+^ glial cells of the CC occurs also from E14.5 **(k)** to E16.5**(l)**. Bar = 60 µm in **a-l**.

Hence, these results indicate that Nkx2.1 expression is only maintained in the astrocyte population of the CC region from E14.5 to E16.5.

### Nkx2.1 controls gliogenesis in embryonic telencephalon

To further investigate the function of Nkx2.1 in regulating embryonic gliogenesis, we performed immunohistochemistry for astroglial (GLAST and GFAP) and polydendroglial (NG2) markers in control and *Nkx2.1^-/-^* embryos expressing inactivated truncated Nkx2.1 (mut-Nkx2.1) at E18.5 (Fig. 4). For control, we made use of both homozygous (*Nkx2.1^+/+^*) and heterozygous (*Nkx2.1^+/^-*) mice. In control mice, GLAST^+^ (n=2) or GFAP^+^ (n=4) astroglia, and NG2^+^ (n=5) polydendrocytes were clearly visible in the CC and its surrounding regions, in the medial cortical area as well as in the septum (Fig. 4a-c and 4e). In contrast, we observed a drastic loss of astroglia (n= 2 stained for GLAST; n=3 stained for GFAP) and polydendrocytes (n=4) in all midline dorsal regions in *Nkx2.1^-/-^* mice (Fig. 4f-h and 4j). However, Nkx2.1 inactivation did not affect the number and organization of radial glia within the GW, in accordance with the absence of Nkx2.1 expression in these glial cell types (compare Fig. 4a,b and 4f,g). Quantitative measurements made with the astrocyte marker, GFAP (Fig. 4k) and polydendrocyte marker, NG2 (Fig. 4l) confirmed the drastic loss (70 to 100%) of astroglia and polydendrocytes in the midline dorsal telencephalic areas in *Nkx2.1^-/-^* mice. There was a drastic disappearance of GFAP^+^ astrocytes in the CC and its surrounding areas (IG and MZG) in the *Nkx2.1^-/-^* embryos (n=3) compared to control embryos (n=4) (p-value=0.0056 for CC, 0.0216 for IG, 0.0067 for MZG, Fig. 4k). Interestingly, analysis with GFAP also revealed the loss of a subpopulation of GFAP^+^ radial glial precursors within the ventral telencephalon, in the mutant POA* and MGE* VZ (Fig. 4i, p-value=0.0496 for MGE and 0.0247 for POA; Fig. 4k). Additionally, there was also a near complete loss (99 to 100%) of NG2^+^ polydendrocytes in the medial cortical areas of *Nkx2.1^-/-^* embryos (n=3) compared to control embryos (n=5) (p-value=0.0334 for CC medial and 0.0191 for CC lateral; Fig. 4j and 4l).

**Figure 4.**
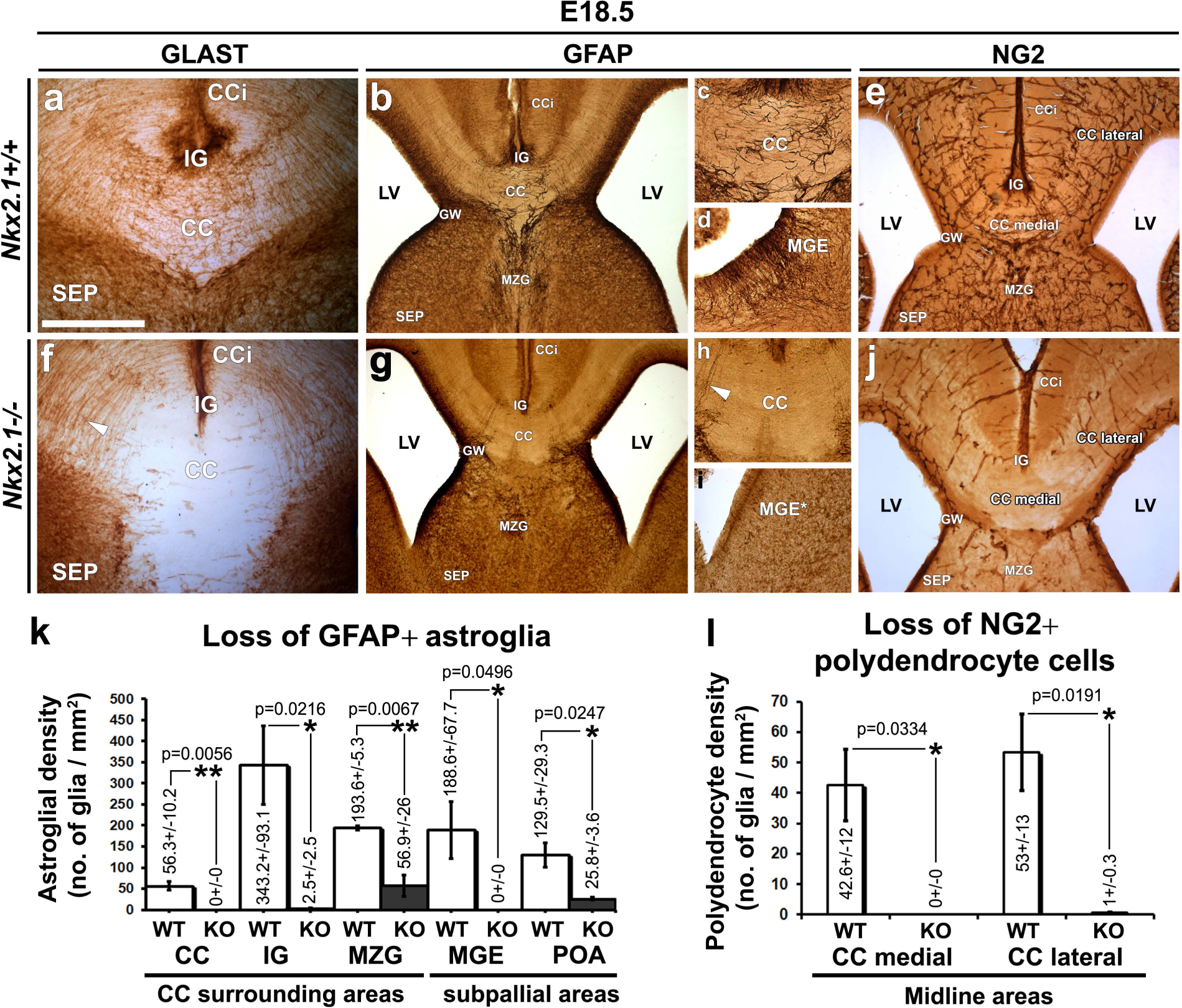
Loss of different glial cell types in the CC, medial cortical areas and subpallium of *Nkx2.1^-/-^* mice brains. DAB staining for GLAST (n=2 for WT and *Nkx2.1^-/-^*) **(a** and **f)**, GFAP (n=4 for WT and 3 for *Nkx2.1^-/-^*) **(b-d** and **g-i)** and NG2 (n=5 for WT and n=4 for *Nkx2.1^-/-^*) **(e** and **j)** on CC and MGE coronal sections from wild-type **(a**-**e)** and *Nkx2.1^-/-^* **(f**-**j)** mice at E18.5. **c** and **h** are higher power views of the CC region seen in **b** and **g**, respectively.**d** and **i** are higher power views of the MGE region. DAB staining for GLAST, GFAP and NG2 revealed a drastic loss of astroglial and polydendroglial cell types from the CC and surrounding areas and from the MGE of the *Nkx2.1^-/-^* mice compared to wild-type mice (compare **f** with **a**, **g-h** with **b-c**, **i** with **d** and **j** with **e**). Only the GFAP^+^ radial glial cells originating from the Nkx2.1^-^ glial wedge (GW) bordering the CC remained unaffected (white arrowhead in **f** and **h**). **(k** and **l)** Bars (mean ± SEM from a sample of n=4 brains in WT and n=3 brains in *Nkx2.1^-/-^* for GFAP and n=5 in WT and n= 3 in *Nkx2.1^-/-^* for NG2) represent the cell densities of GFAP^+^ or NG2^+^ glial cells/mm^2^. The quantification of the GFAP^+^ (p-value=0.056 for CC, 0.0216 for IG, 0.0067 for MZG, 0.0496 for MGE, and 0.0247 for POA) and NG2^+^ (p-value=0.0334 for CC medial and 0.0191 for CC lateral) glial cell density showed a drastic and significant loss of these cells in the CC and surrounding areas as well as in medial cortical areas of the *Nkx2.1^-/-^* brain compared to the wild-type brains. **(CC)** corpus callosum; **(CCi)** cingulate cortex; **(GW)** glial wedge; **(IG)** induseum griseum; **(LV)** lateral ventricle; **(MZG)** midline zipper glia;**(MGE)** medial ganglionic eminence; **(SEP)** septum. Bar = 500 µm in **b**, **e**, **g** and **j**; 250 µm in **a**, **f**, **c**, **d**, **h** and **i**.

Furthermore, since we identified different *Nkx2.1*-derived glial cell populations, both astrocyte-like and polydendrocyte-like, this prompted us to investigate in further the function of Nkx2.1 in glia specification while considering these different glial cell types (Figure 2). To this purpose, we performed immunostaining for Olig2 and GLAST on telencephalic CC sections from both controls (n=4) and *Nkx2.1^-/-^* (n=2) mice. In the *Nkx2.1^-/-^* CC, a significant reduction (around 70%) in the cell density of GLAST^+^/Olig2^-^ astrocytes was observed when compared to the WT (p-value= 0.0297), while no differences were detected for GLAST^+^/Olig2^+^ astroglial cell densities (p-value= 0.4469) (Fig. 2a-e). The GLAST/Olig2^+^/ polydendrocyte population was nearly completely lost (around 98%) in the CC (p-value= 0.0242; Fig. 4e). Hence, these results show that Nkx2.1 regulates both the GLAST^+^/Olig2^−^ astroglial and GLAST^−^/Olig2^+^ polydendroglial populations but not the GLAST^+^/Olig2^+^ astroglial population (also illustrated in Figure 1-figure supplement 3).

In order to exclude the possibility of cell death being the central reason for the marked decrease in the number of glial cells we observed, we analyzed the controls (n=4 for CC region; n= 5 for POA region) and *Nkx2.1^-/-^* (n=6 for CC region; n=10 for POA region) brains at E16.5 for cleaved-caspase 3, a key biomarker for apoptosis (Figure 4-figure supplement 1a-d and 1i). We also performed terminal deoxynucleotidyl transferase-mediated dUTP-biotin nick end labeling (TUNEL) assay which detects DNA fragmentation that results from different cell death processes (n=16 for CC in controls, n=22 for CC in knockouts; n=6 for POA in controls, n=5 for POA in knockouts; n=10 for MGE in controls, n= 11 for MGE in knockouts; n=7 for SEP in controls, n=14 for SEP in knockouts; Figure 4-figure supplement 1e-h and 1j). The DNA binding dye, Hoechst was also used to visualize the pyknotic nuclei and in addition, to determine if there are any differences in nuclear size or nuclear morphology between the WT and mutant brains. The quantification of the absolute number of dying cells, labeled by the cleaved-caspase 3 (Figure 4-figure supplement 1i), and by TUNEL staining (Figure 4-figure supplement 1j), revealed no significant differences between the *Nkx2.1^-/-^* brains and the control brains in any of the observed telencephalic regions namely the CC, the MGE, the POA or the septum (p-value= 0.1225 for CC and 0.4618 for POA with cleaved caspase 3 staining; p-value= 0.7934 for CC, 0.8193 for POA, 0.4032 for MGE, and 0.4879 for SEP with TUNEL). The size and morphology of the cell nuclei was comparable in both WT and mutant brains, with no significant differences observed.

These results show that the glial cells occupying the CC are under the regulation of Nkx2.1. Also, the observed loss of astrocytes and polydendrocytes in the *Nkx2.1^-/-^* telencephalon is not due to the glial cell death.

### Nkx2.1 regulates the proliferation of astrocyte ventral progenitors in embryonic brains

The loss of specified glia in *Nkx2.1^-/-^* mice may be owing to insufficient proliferation of ventral glial Nkx2.1-precursors of the progenitor zones, namely MGE, AEP/POA and TS. To investigate further, we performed double immunohistochemical staining for Nkx.2.1 and for the radial glia/astrocytic marker, GLAST on coronal sections of the precursor regions in both *Nkx2.1^+/+^* or *Nkx2.1^+/^-* control (n=4) and *Nkx2.1^-/-^*(n=4) mice brains at E16.5. Likely both, the mutated Nkx2.1 (mut-Nkx2.1) and the WT Nkx2.1 proteins are similarly recognized by the anti-Nkx2.1 antibody indicating that the mutated protein conserves an intact epitope sufficient for recognition by the antibody (Corbin et al., 2003).

In the VZ, the SVZ and the mantle zone of the control MGE, many GLAST^+^ precursors and differentiated astroglia co-expressed Nkx2.1 (Fig. 5a-c and 5h, solid arrowheads) whereas in the germinal and mantle zones of the mutant MGE* of *Nkx2.1^-/-^* mice, only few GLAST^+^ precursors and astroglia co-labeled for the mut-Nkx2.1 were observed (Fig. 5d-f, solid arrowheads). This observed difference may be attributed to the fact, as shown before, that although a MGE-like structure forms in the mutant (called as MGE*), it has been re-specified to a more dorsal LGE-like fate (Sussel et al., 1999). Similar reduction of cells labeled for GLAST and the mut-Nkx2.1 were seen in the mutant POA* of *Nkx2.1^-/-^* mice (Fig. 5i-j). The quantitative analyses revealed a drastic and significant decrease of the total cells (50 to 85%) and GLAST^+^ precursors (45 to 86%) expressing the mutated Nkx2.1 in the VZ, SVZ of MGE*, POA* and TS* (p-value < 0.0001 in the VZ of MGE, POA and TS in Fig. 5k-l and p-value=0.0139 for the SVZ of MGE in Fig. 5k). Consequently, the number of GLAST^+^ differentiated astrocytes co-expressing the mut-Nkx2.1 in the parenchyma (striatum, LPOA/LH, septum) of the *Nkx2.1^-/-^* (n=4) was severely decreased (60 to 80%) as compared to control mice (n=4) (p-value<0.0001 for striatum, LPOA and septum in Fig. 5l). Thus, these results indicate that the mutation of Nkx2.1 results in the severe loss of precursors in the mutant mice brains.

**Figure 5.**
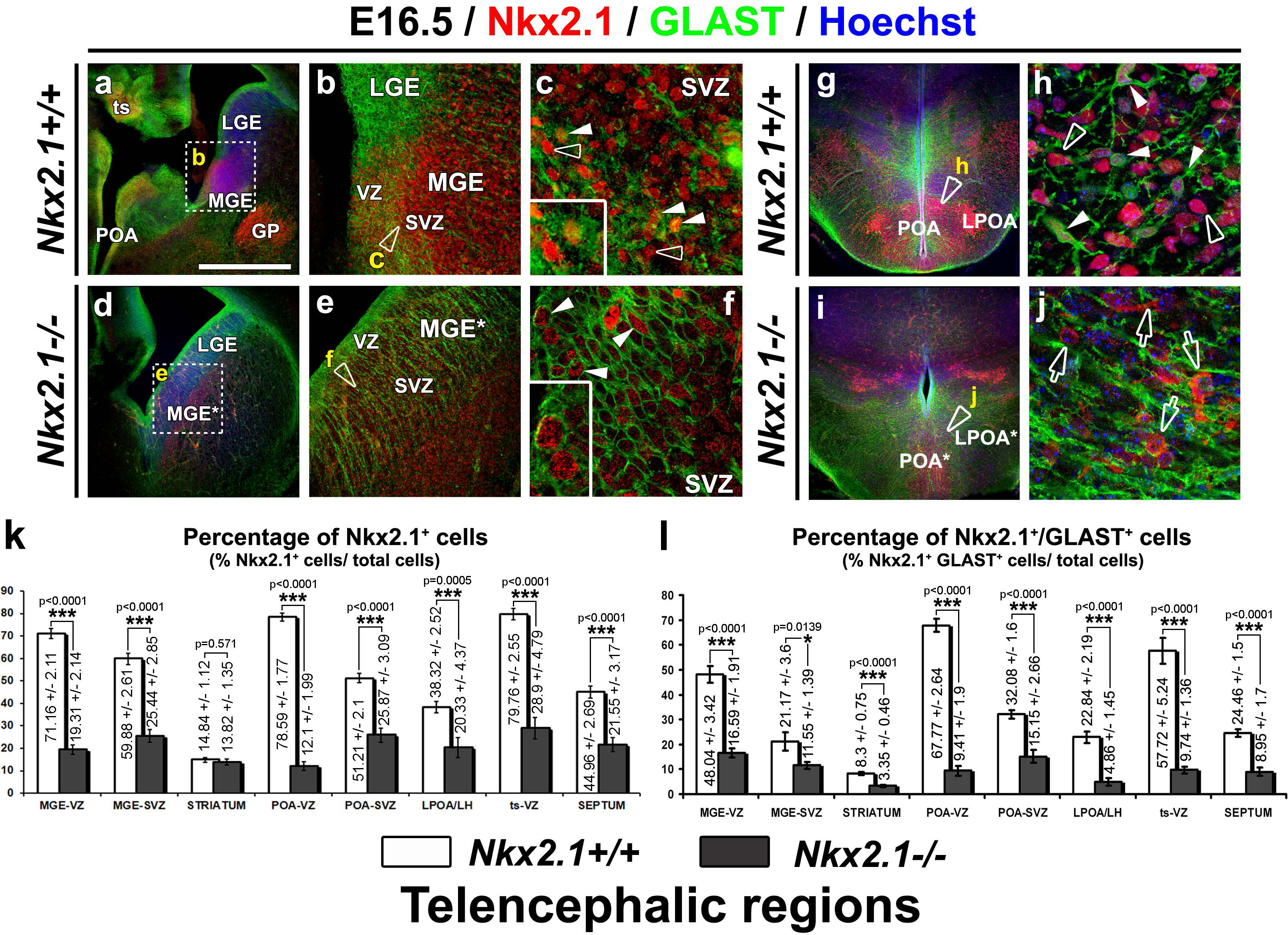
Incapacity of precursors to generate astrocytes in *Nkx2.1*^-/-^ mice brains. **(a-c** and **g-h)** Double immunohistochemical staining for Nkx2.1 and GLAST on MGE **(a-c)** and POA **(g-h)** coronal sections from wild-type (n=4) mice brains at E16.5. **(d-f** and **i-j)** Double immunohistochemical staining for mutated Nkx2.1 (mut-Nkx2.1) and GLAST on MGE* **(d-f)** and POA* **(i-j)** coronal sections from *Nkx2.1^-/-^* (n=4) mice brains at E16.5. Cell nuclei were counterstained in blue with Hoechst **(a, d, g-h and i-j)**. **b, c, e, f, h** and **j** are higher power views of the regions seen in **a, d**, **g** and **i** respectively. **(a-c)** In the germinal regions of the wild-type MGE, numerous Nkx2.1^+^ progenitors were GLAST^+^ (solid arrowheads and inset in **c**), while some other were GLAST^-^ (open arrowheads in **c**). **(d-f)** In *Nkx2.1^-/-^* MGE* germinal regions, only few GLAST^+^ progenitors expressed the mutated Nkx2.1 protein (solid arrowheads and inset in **f**). **(g-h)** In the parenchyma of wild-type POA, many GLAST^+^ astroglial cells (solid arrowheads in **h**) and neurons expressed Nkx2.1. **(i-j)** In the parenchyma of *Nkx2.1^-/-^* POA*, GLAST^+^ astroglial cells have disappeared and only few neurons expressing the mutated Nkx2.1 protein are observed (open arrows in **j**). **(GP)** globus pallidus; **(LGE)** lateral ganglionic eminance; **(LPOA)** lateral POA; **(MGE)** medial ganglionic eminence; **(MGE*)** mutant medial ganglionic eminence; **(POA)** preoptic area; **(POA*)** mutant preoptic area; **(SEP)** septum; **(SVZ)** subventricular zone; **(TS)** triangular septal nucleus, **(VZ)** ventricular zone. Bar: 100 µm in **b** and **e**; 45 µm in **c** and **f** and 50 µm in **h** and **j**. **(k-l)** Bars (mean ± SEM from n=4 brains in WT and n=4 brains in Nkx2.1^-/-^ mice) represent the percentage of Nkx2.1^+^ cells **(k)** and of Nkx2.1^+^/GLAST^+^ precursors and astroglial cells **(l)** in WT (white columns) and *Nkx2.1^-/-^* (black columns) subpallial germinal (MGE, POA and TS: VZ and SVZ) and parenchymal (striatum, LPOA/LH, septum) telencephalic regions at E16.5. **(k** and **l)** The number of cells **(k)** and GLAST^+^ **(l)** precursors and post-mitotic cells expressing the mutated Nkx2.1 was drastically decreased in all the subpallial telencephalic regions of the *Nkx2.1^-/-^*. The p-values are indicated above the respective graphs within the figure panels k to l.

To ascertain the cell proliferation status of precursors at the germinal zones, we made use of the S-phase marker, BrdU. The rate of cell proliferation was studied in *Nkx2.1^+/+^* (n=8) or *Nkx2.1^+/^-* (n=3) control and *Nkx2.1^-/-^* (n=8) mice with a principal attention at E16.5 when the bulk of embryonic telencephalic glia is generated. In the control brains, both in the VZ and SVZ of the MGE, numerous Nkx2.1^+^ precursors were co-labeled with BrdU (Fig. 6a-c, solid arrowheads). For quantification, we used n=4 control and n=4 knockouts (Fig. 6g-j). In the mutant MGE* of *Nkx2.1^-/-^* mice, we observed a significant reduction in the BrdU^+^ progenitors of the VZ and SVZ compared to the control MGE (compare Fig. 6c to 6f; p-value < 0.0006 in the VZ of MG and p-value <0.0064 in the SVZ of MGE in Fig. 6g). Intriguingly, an increase in number of total BrdU^+^ cells was seen in the POA* VZ and SVZ regions of the *Nkx2.1^-/-^*. Nonetheless, remaining BrdU^+^ precursors only very rarely expressed the mutated Nkx2.1 in both the mutant MGE* and POA* regions (Fig. 6h). The number of dividing cells labeled for the mut-Nkx2.1 was severely reduced (75 to 80%) in the *Nkx2.1^-/-^* brains (p-value <0.0001 in the VZ of MGE, POA and in the SVZ of MGE, POA in Fig. 6h). The mutation of Nkx2.1 results in the incapacity of Nkx2.1-derived precursors to divide (p-value <0.0001 in the VZ and SVZ of MGE and p-value <0.0005 in the SVZ of POA in Fig. 6i). By contrast, dividing cells that did not express the mut-Nkx2.1 in the *Nkx2.1^-/-^* brains were either not affected or up-regulated in the mutant MGE* and POA* regions (Fig. 6j).

**Figure 6.**
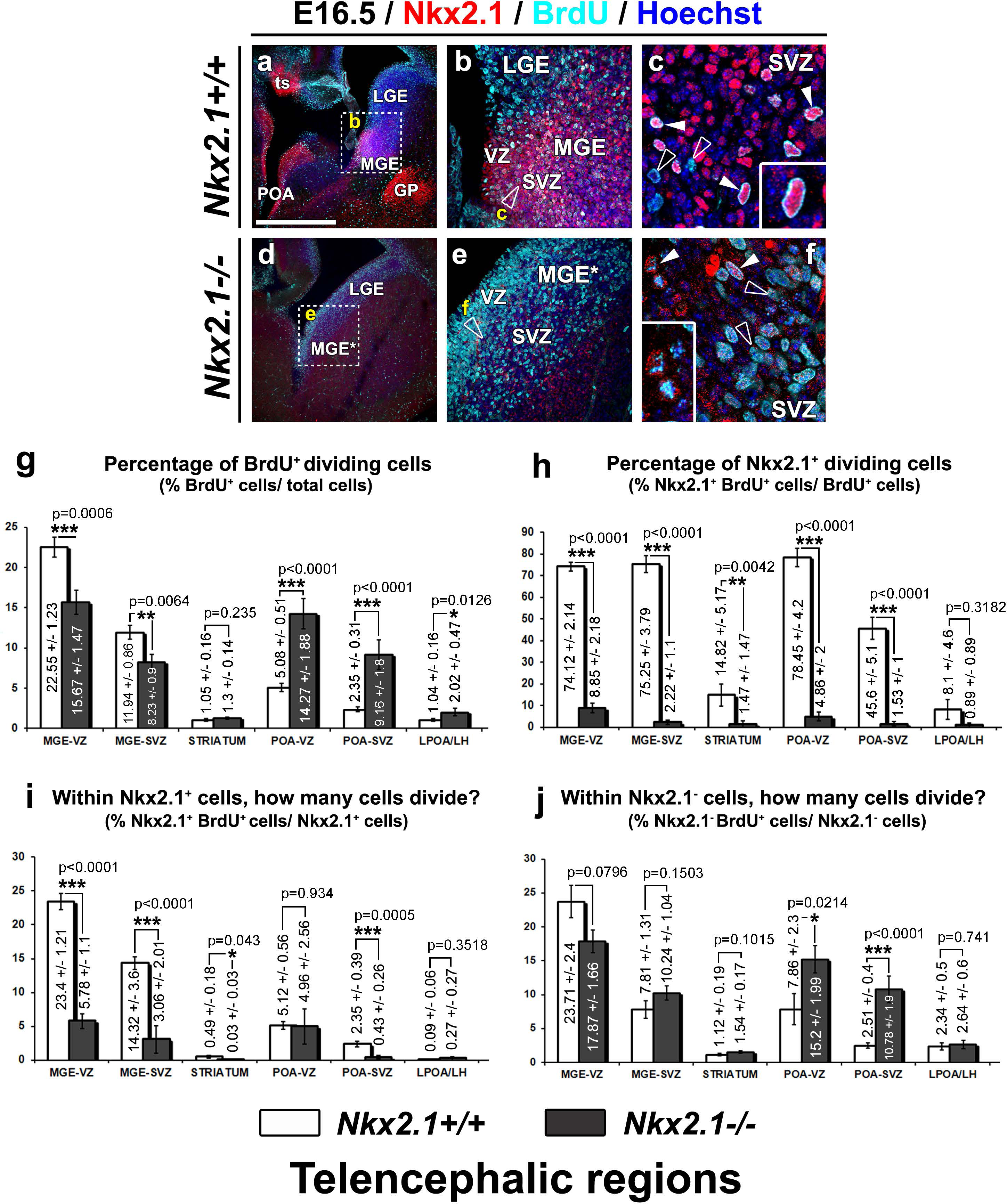
Incapacity of Nkx2.1^+^ precursors to divide after Nkx2.1 inactivation. **(a-c)** Double immunohistochemical staining for Nkx2.1 and 5-bromo-2'- deoxyuridine (BrdU) on telencephalic coronal sections from wild-type (n=4) mice brains at E16.5. **(d-f)** Double immunohistochemical staining for mutated Nkx2.1 and BrdU on telencephalic coronal sections from *Nkx2.1^-/-^* (n=4) mice brains at E16.5. Cell nuclei were counterstained in blue with Hoechst **(a-f)**. **b** and **e** are higher power views of the MGE squared regions seen in **a** and **d**. **c** and **f** are higher magnifications of the MGE seen in **b** and **e**, respectively. **(a-c)** In the VZ and the SVZ of the wild-type MGE, AEP/POA and TS, numerous Nkx2.1^+^ precursors (in red) co-labelled for BrdU (in light blue) were dividing at E16.5 (solid arrowheads and inset in **c**). Other BrdU^+^ dividing cells were not labelled by Nkx2.1 (open arrowheads in **c**). **(d-f)** In the VZ and the SVZ of the *Nkx2.1^-/-^* MGE (MGE*), numerous precursors co-labelled by the BrdU were also seen to divide (open arrowheads in **f**), but only few dividing cells were expressing the mutated Nkx2.1 protein (solid arrowheads and inset in **f**). **(LGE)** lateral ganglionic eminance; **(MGE)** medial ganglionic eminence; **(MGE*)** mutant medial ganglionic eminence;**(POA)** preoptic area; **(SVZ)** subventricular zone; **(TS)** triangular septal nucleus, **(VZ)** ventricular zone. Bar = 675 µm in **a** and **d**; 100 µm in **b** and **c** and 45 µm in **c** and **d**. **(g-j)** Bars (mean ± SEM from n=4 brains from WT and n=4 brains from Nkx2.1^-/-^ mice) represent the percentage of the BrdU^+^ dividing cells **(g)**; the percentage of BrdU^+^dividing cells which are also positive for Nkx2.1 or mutated Nkx2.1 **(h)**; the percentage of cells for Nkx2.1 or mutated Nkx2.1 that divided **(i),** and the percentage of Nkx2.1^-^ cells that divided **(j)** in WT and *Nkx2.1^-/-^* germinal (MGE, POA and TS: VZ and SVZ) and parenchymal (striatum, LPOA/LH) telencephalic regions at E16.5. **(g)** A significant decrease of the BrdU^+^ dividing precursors in the VZ and SVZ of the MGE* was balanced by a significant increase of BrdU^+^ dividing precursors in the VZ, SVZ of the POA* of Nkx2.1^-/-^; **(h)** a drastic and significant decrease in the dividing cells which expressed mutated Nkx2.1 was observed in all regions of Nkx2.1^-/-^; **(i)** the cells that express the mutated Nkx2.1, lost their capacity to divide in the VZ and SVZ of the MGE*, the striatum and the SVZ of the POA*; **(j)** by contrast, the cells that do not express mutated Nkx2.1, still divided normally in the MGE* and maintained the capacity to divide in the VZ and SVZ of the POA* of Nkx2.1^-/-^. The p-values are indicated above the respective graphs within the figure panels g to j.

Altogether, these observations indicate that the transcription factor Nkx2.1 controls the proliferation step of the original Nkx2.1^+^ precursors in the MGE and the POA, the subpallial domains that majorly generate early embryonic astroglia.

### Nkx2.1 regulates the differentiation of Nkx2.1-derived astroglia in embryonic brains

Next, we aimed to analyze the specification of Nkx2.1^+^ precursors capable of generating early astroglial cells by observing the MGE- and the POA-derived neurosphere differentiation at E14.5. After 7 days *in vitro* (DIV), control MGE and POA neurospheres were able to differentiate and generate two cell types of the brain (Arsenijevic et al., 2001) which were GFAP^+^ mature astrocytes (Fig. 7a-b) and ßIII-tubulin^+^ post-mitotic neurons (not shown). GFAP^+^ astrocytes were observed to be uniformly dispersed on the entire surface of the spheres (Fig. 7a-b). Using immunohistochemistry for Nkx2.1, we found that in the neurospheres derived from control MGE and POA, Nkx2.1 was expressed in the nucleus of about 50% of the GFAP^+^ astroglia (Fig. 7b and e; solid arrowheads). Thereafter, *in vitro* differentiation of E14.5 *Nkx2.1^-/-^* mutant MGE* and POA*-derived neurospheres revealed that, precursors expressing the mut-Nkx2.1 stopped to differentiate into mut-Nkx2.1^+^/GFAP^+^ astroglia (Fig. 7c-d). Quantification showed that the mutation of Nkx2.1 in the MGE* and POA* neurospheres of *Nkx2.1^-/-^* induced a significant decrease of mut-Nkx2.1^+^/GFAP^+^ astroglia (p-value <0.0001 in the MGE and POA neurospheres in Fig. 7e). While more than 50% of differentiated GFAP^+^ astroglia expressed Nkx2.1 in control neurospheres, less than 10% of GFAP^+^ astroglia expressed mut-Nkx2.1 in mutant neurospheres (Fig. 7e). It indicates that in mutant MGE* and POA* neurospheres, mut-Nkx2.1^+^ precursors have nearly completely lost the capacity to differentiate into astroglia.

**Figure 7.**
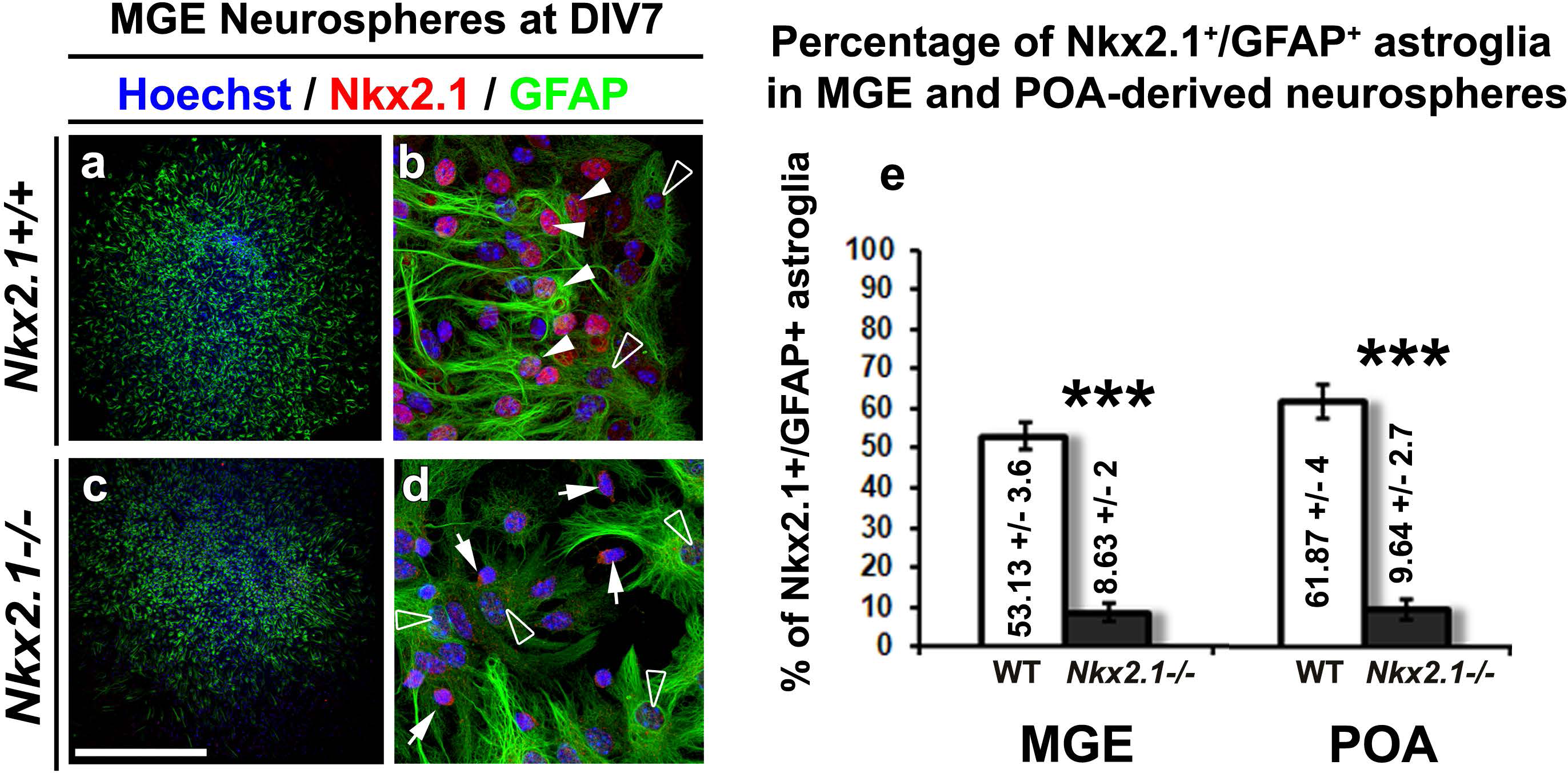
Nkx2.1^+^ MGE and POA stem cells do not differentiate into astrocytes after *Nkx2.1* inactivation. **(a-b)** Double immunocytochemistry for Nkx2.1 and GFAP on MGE-derived neurospheres from E14.5 wild-type mice (n=8) after 7 days *in vitro*(DIV). **(c-d)** Double immunocytochemistry for mutated Nkx2.1 and GFAP on MGE*-derived neurospheres from E14.5 *Nkx2.1^-/-^* mice (n=6) after 7 days *in vitro*(DIV). Cell nuclei were counterstained in blue with Hoechst **(a-d)**. **b** and **d** are higher power views of the regions seen in **a** and **c** respectively. In neurospheres derived from wild-type MGE, numerous GFAP^+^ astrocytes were labelled for Nkx2.1 (solid arrowheads in **b**), while some others were not (open arrowheads in **b**). By contrast, in neurospheres derived from *Nkx2.1^-/-^* MGE*, the GFAP^+^ astrocytes were never observed to be co-labelled for the mutated Nkx2.1 (open arrowheads in **d**), but neuronal cells still express low levels of the mutated Nkx2.1 protein (arrows in **d**). Bar = 675 µm in **a** and **c** and 50 µm in **b** and **d**. **(e)** Bars (mean ± SEM from a sample of n=21 MGE, n=34 POA WT neurospheres and n=22 MGE*, n=28 POA* *Nkx2.1^-/-^* neurospheres) represent the percentage of GFAP^+^ astrocytes labeled for Nkx2.1 in MGE– or POA–derived neurospheres from wild-type and of GFAP^+^ astrocytes labeled for mutated Nkx2.1 in MGE*– or POA*– derived neurospheres form *Nkx2.1^-/-^* mice brains. Neurospheres originating from *Nkx2.1^-/-^* MGE* and POA* nearly lost the capacity to produce Nkx2.1-derived astrocytes (p-value<0.0001 for both).

### Nkx2.1 directly regulates the expression of the GFAP astroglial regulatory gene

It is clear from our above mentioned results that the astroglial cell populations of the embryonic telencephalon are derived from Nkx2.1^+^ progenitors, and Nkx2.1 regulates the astrocyte precursor cells proliferation and differentiation. In order to ascertain if the transcription factor Nkx2.1 regulates expression by binding onto the promoter sequences of the astroglial GFAP regulatory gene in the brain, we performed chromatin immunoprecipitation assay on lysates of E16.5 embryonic brains. Firstly, we searched for DNA elements matching the consensus NK2 family binding sequence [GNNCACT(T/C)AAGT(A/G)(G/C)TT] (Guazzi et al., 1990) in the upstream promoter regions of the regulatory gene GFAP and of the negative control gene Ngn2 that regulates dorsal precursors (Fode et al., 2000). In the absence of the complete consensus binding sequence, the core binding sequence T(C/T)AAG was chosen for analysis. As a positive control, we included Lhx6 in our analysis for which the site for the binding of Nkx2.1 is already known (Du et al., 2008). Secondly, after shortlisting the position of putative Nkx2.1 binding sites, primers for the sequence flanking all the shortlisted binding sites (up to three) were made. We then performed the PCR on the crosslinked and sonicated DNA pulled down using an anti-Nkx2.1 monoclonal antibody. The amplification of a putative Nkx2.1 binding sequence located in the PCR product within the astroglial gene for GFAP (Fig. 8a) was found to be positive in the brain samples chromatin immunoprecipitated with the Nkx2.1 antibody, however, as expected no positive interaction was detected for Ngn2 (Fig. 8c). Also, the amplification of the PCR product comprising the already known Nkx2.1 binding site within the Lhx6 promoter was positive upon immunoprecipitation with the anti-Nkx2.1 antibody (Fig. 8b). Furthermore, the PCR fragment(s) amplified for the GFAP promoter contained the core sequence CTCAAGT of the Nkx2.1 consensus binding sequence. Thus, these results suggest that, *in vivo*, Nkx2.1 binds the promoter region of the GFAP astroglial regulatory gene, which contains the aforementioned highly conserved core-binding sequence of the consensus binding site.

**Figure 8.**
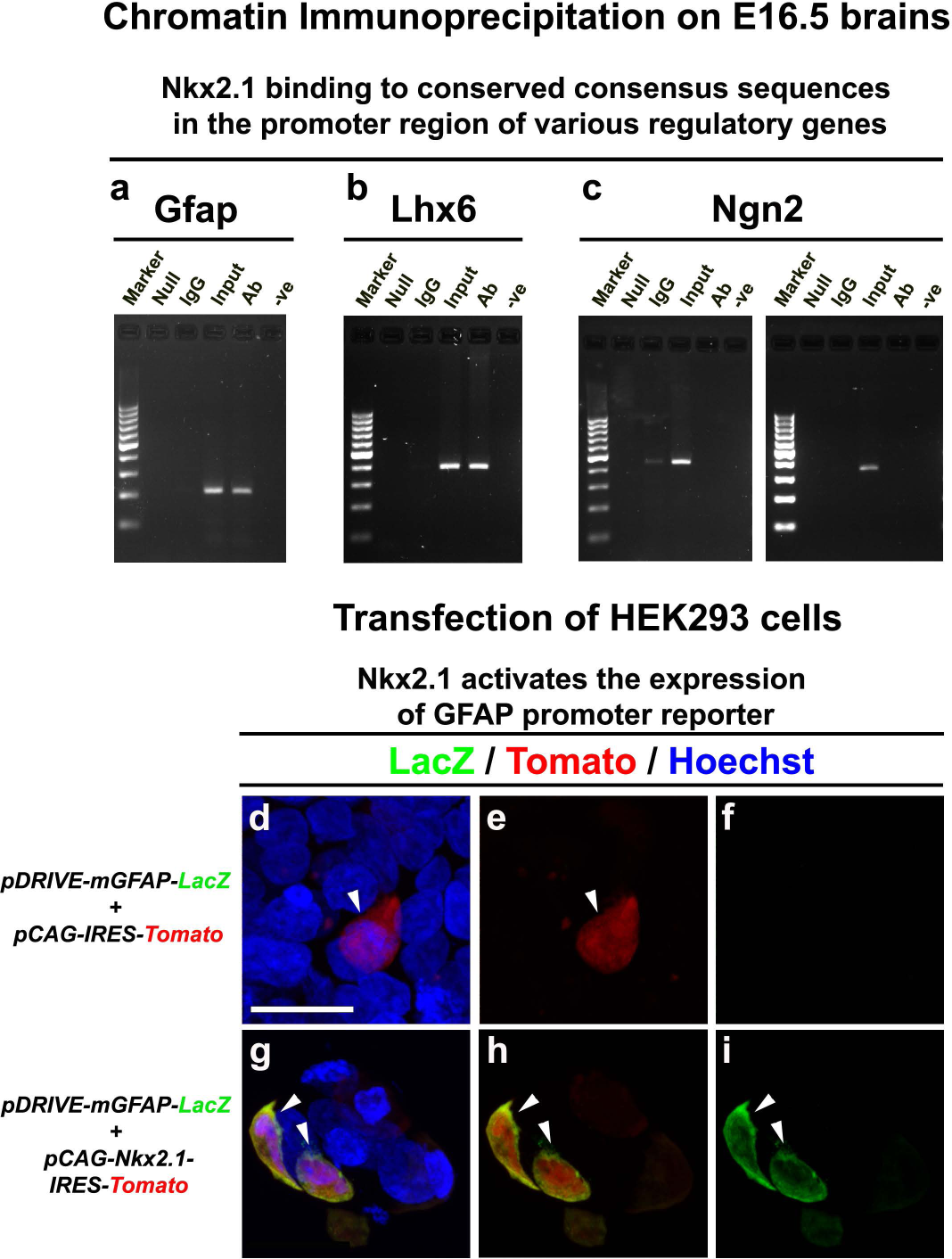
Binding of Nkx2.1 to conserved binding sequences in the promoters of various glial regulatory genes. Amplification of the Nkx2.1 putative binding sequences located in a 206 bp PCR product within the GFAP promoter **(a)** and in a 391 bp PCR fragment within the Lhx6 promoter **(b)** after chromatin immunoprecipitation with the Nkx2.1 antibody. Input DNA was added as the positive loading control as it contains the crosslinked sonicated genomic DNA taken before chromatin immunoprecipitation with the Nkx2.1 antibody, and a strong signal was observed for all the promoter regions. No amplification of the Nkx2.1 core sequence (tcaag) located in two, 410 bp and 352 bp, PCR products within the Ngn2 promoter was detected **(c)**. No signal was detected in the null control, wherein no antibody was added for chromatin immunoprecipitation and in the negative control, wherein no DNA was added while performing the PCR **(a-c)**. A very faint signal was detected in some of the samples immunoprecipitated with non-specific control IgG but its intensity was much lower than the intensity of the input DNA and the test DNA (containing promoter region). These results suggest that Nkx2.1 binds the promoter regions of various glial regulatory genes at a conserved Nkx2.1 binding sequence *in vivo*. The figure represents one of the three independently performed assays. Identical results were also obtained for the same glial regulatory genes in the E14.5 brain samples. As a control, the human embryonic kidney 293 (HEK293) cells were co-transfected with 2.0 µg of two reporter constructs, namely, the *pDRIVE-mGFAP-LacZ* expression plasmid containing the LacZ reporter under the control of the mouse 1679 bp upstream GFAP promoter sequence, and the *pCAG-IRES-Tomato* plasmid constitutively expressing the Tomato reporter under the control of the pCAG promoter **(d-f)**. To test the binding of the Nkx2.1 to the GFAP promoter sequence, the HEK293 cells were co-transfected with 2.0 µg of two reporter constructs, namely, the *pDRIVE-mGFAP-LacZ* expression plasmid, and the *pCAG-Nkx2.1-IRES-Tomato* plasmid expressing Nkx2.1 protein tagged with Tomato under the control of the constitutive promoter pCAG **(g-i)**. Cell nuclei were counterstained in blue with Hoechst. Activation of the LacZ reporter was seen upon addition of the Nkx2.1 expression vector, thus confirming that Nkx2.1 activates the GFAP promoter, most probably through the binding sequence that we identified. Bar = 50 µm.

Thereafter, to confirm the direct influence of the binding of Nkx2.1 to this upstream binding sequence towards the transcription of GFAP astroglial regulatory gene, we performed co-transfection studies in the HEK293 cells. We made use of the expression plasmid *pDRIVE-mGFAP*, which contains the LacZ reporter under the control of the mouse upstream −1679 bp GFAP promoter sequence (and includes the putative Nkx2.1 binding site identified above), and the *pCAG-Nkx2.1-IRES-Tomato* plasmid constitutively over-expressing the Nkx2.1 protein and the Tomato protein under the control of the *pCAG* promoter. Co-transfection of both plasmids in the HEK293 cells resulted in robust expression of the LacZ reporter (Fig. 8g-i). Contrastingly, almost no expression was apparent upon transfection of the *pDRIVE-mGFAP* plasmid with the control *pCAG-IRES-Tomato* plasmid lacking the *Nkx2.1* cDNA (Fig. 8d-f). Thus, these results suggest that the activation of the *GFAP* promoter fragment requires the presence of Nkx2.1, which most probably recognizes the binding sequence identified by us.

These results, altogether, show that the *Nkx2.1* homeobox gene indeed regulates proliferation and differentiation of the astroglia that occupy the CC region during late embryonic ages.

## Discussion

Nkx2.1 has been implicated during the specification of GABAergic interneurons and oligodendrocytes that occupy the embryonic telecephalon (Anderson et al., 2001, Corbin et al., 2001, Kessaris et al., 2006, Kimura et al., 1996, Marin and Rubenstein, 2001, Sussel et al., 1999). Recently, our group has shown that Nkx2.1 controls the production of GABAergic interneurons, astrocytes and polydendrocytes that populate the embryonic ventral telencephalon (Minocha et al., 2015a, Minocha et al., 2015b). In this study, our results unravel that Nkx2.1 also regulates the generation of dorsal astroglia that populate the corpus callosum and its surrounding regions during late embryonic stages. Nkx2.1 mediates its control over astroglia through regulation of both proliferation and differentiation of Nkx2.1^+^ precursors present in the ventral progenitor regions, namely the MGE, the AEP/POA and the TS (Minocha et al., 2015a, Minocha et al., 2015b). By controlling the production of neuron and glia that populate the entire telencephalon, Nkx2.1 is a key factor for brain shaping during embryonic development.

### Multilevel regulation of embryonic astrogliogenesis by Nkx2.1

The Nkx2.1-derived cell population in embryos is broadly divided into astrocyte-like and polydendrocyte-like based on their expression profile, and is generated maximally between E14.5-to-E16.5. Only Nkx2.1^+^ astrocyte-like cells that are GLAST^+^ and/or GFAP^+^ continue to maintain Nkx2.1 expression while other Nkx2.1-derived cells that are NG2^+^/Olig2^+^ polydendrocyte-like no longer express Nkx2.1 as soon they differentiate. Loss of Nkx2.1 function in *Nkx2.1^-/-^* mice leads to a drastic reduction in the number of both astroglia and polydendrocytes in the midline dorsal (CC, IG, MZG) and within the ventral telencephalic (mutant MGE* and POA*, and septum) regions. The loss of astroglia and polydendrocytes is not accompanied with a concomitant increase in apoptotic cells. Hence, the results indicated that the loss might be due to incapacity of the precursors to generate astroglia and polydendrocytes. Indeed, further analyses revealed that the loss of glia is accompanied with a decrease of *Nkx2.1*-derived precursor division capacity and astroglial differentiation. Accordingly, we observed a drastic decrease in presence of total cells and GLAST^+^ precursors expressing the mut-Nkx2.1 in the VZ, SVZ of mutant MGE*, mutant POA* and the TS region, and of the GLAST^+^ differentiated astrocytes expressing the mut-Nkx2.1 in the parenchyma (striatum, LPOA/LH, septum) of the *Nkx2.1^-/-^* compared to the WT mice. The decreased presence of precursors and differentiated astroglial population was accompanied with reduced proliferative status of the BrdU^+^ dividing cells labeled for mut-Nkx2.1 in VZ and SVZ of mutant MGE*, mutant POA* and the septal nucleus region of *Nkx2.1^-/-^* mice compared to the WT precursors. Additionally, *in vitro* differentiation of E14.5 *Nkx2.1^-/-^* mutant MGE*, POA*-derived neurospheres revealed that, after Nkx2.1 inactivation, the progenitors were unable to differentiate into GFAP^+^ astroglia expressing the mut-Nkx2.1^+^ significantly though they still retained the capacity to generate post-mitotic neurons. Hence, the reduction in number of astroglia and polydendrocytes can be attributed to a deficit in proper proliferation and differentiation of precursors in three subpallial domains in the absence of Nkx2.1. Chromatin immunopreciptation analysis also suggests that the Nkx2.1 might mediate this control by direct activation of astroglial gene promoter region, GFAP here.

Thus, Nkx2.1 regulates the generation and specification of dorsal telencephalic astroglia through a multilevel control that involves (i) control over proliferation of Nkx2.1^+^ precursors, (ii) regulation of differentiation of Nkx2.1^+^ precursors, and lastly, (iii) transcriptional control over astroglial gene like GFAP.

### Nkx2.1 is important for several aspects of proper brain development during embryogenesis

Previous reports from our and other groups have shown that not only is Nkx2.1 important for regional specification of the ventral telencephalic regions, MGE and POA, it is also essential for generation of a wide spectrum of Nkx2.1-derived lineages including GABAergic interneurons, polydendrocytes and astrocytes that populate both the dorsal and ventral telencephalon beginning from E12.5 (Anderson, 2001, Corbin et al., 2001, Du et al., 2008, Kessaris et al., 2006, Kessaris et al., 2008, Kimura et al., 1996, Marin et al., 2000, Marin and Rubenstein, 2001, Marin et al., 2010, Minocha et al., 2015a, Minocha et al., 2015b, Nobrega-Pereira et al., 2008, Sussel et al., 1999, Xu et al., 2008). Maximal generation of Nkx2.1-derived cell types occurs around E14.5-to-E16.5, a period characterized by several key developmental events, including midline fusion, and bordering the formation of corpus callosum and anterior commissure and also blood vessel network (Adams and Alitalo, 2007, Larrivee et al., 2009, Minocha et al., 2015a, Minocha et al., 2015b, Paul et al., 2007, Richards et al., 2004). Loss of Nkx2.1 leads to ventral-to-dorsal transformation of the pallidum (Sussel et al., 1999), together with drastic reduction in Nkx2.1-derived cell population leading to structural abnormalities in anterior commissure and blood vessel network (Minocha et al., 2015a, Minocha et al., 2015b). Also, the reduced GABAergic neuronal localization in the Nkx2.1^−/–^ mice lead to callosal axon branching and outgrowth defects in the CC tract (Niquille et al., 2013).

This study shows that Nkx2.1 is able to perform its vast range of roles through regulation of both proliferation and differentiation of Nkx2.1^+^ precursors. It appears that Nkx2.1 mediates some (or all) of these effects through transcriptional regulation of target genes, such as astroglial gene GFAP characterized in this study.

Previous reports have shown that Nkx2.1 regulates the transcription of many genes of the thyroid (Guazzi et al., 1990, Lazzaro et al., 1991, Sussel et al., 1999) and activates pulmonary-surfactant (Boggaram, 2009), as well as pituitary gland genes (Hamdan et al., 1998). Moreover, it has been shown *in vitro*, that Nestin might be a target of *Nkx2.1* (Lonigro et al., 2001). Another group has also seen transcriptional patterns of regulation by Nkx2.1 in early (E11.5) and late (E19.5) mouse lung development (Tagne et al., 2012). Interestingly, in mouse lungs, Nkx2.1 also directly regulates the cell cycle effectors and its loss alters cell cycle progression (Tagne et al., 2012). In the ventral telencephalon, loss of Nkx2.1 function also affects the proliferation of precursors expressing the mut-Nkx2.1. Hence, it is probable that Nkx2.1 displays some functional conservation in brain, thyroid, pituitary, and lung — the four Nkx2.1^+^ identified tissues.

Complex cellular and molecular interactions between glia, neurons and guidance cues produced by them govern the formation of the midline structures such as corpus callosum and anterior commissure. Several Nkx2.1-derived glial and neuronal populations populate these aforementioned structures, and further understanding of the mode of regulation mediated by Nkx2.1 can help better understand the formation of dorsal and ventral telencephalic regions.

## Methods

### Animals

All studies on mice of either sex have been performed in compliance with the national and international guidelines. For staging of embryos, midday of the day of vaginal plug formation was considered as embryonic day 0.5 (E0.5). Wild-type mice maintained in a CD-1/SWISS genetic background were used for developmental analysis of the CC. We used wild-type (+/+) and homozygous mutant *Nkx2.1* mice (Flames et al., 2007, Kimura et al., 1996, Sussel et al., 1999), which are referred as *Nkx2.1 ^+/+^* and *Nkx2.1^-/-^* in this work. We used heterozygous *GAD67*-*GFP* knock-in mice, described in this work as *Gad1-EGFP* knock-in mice (Tamamaki et al., 2003). *Gad1-EGFP* knock-in embryos could be recognized by their GFP fluorescence. PCR genotyping of these lines was performed as described previously (Niquille et al., 2009). We used *GLAST-Cre ERT^TM^*(purchased from Jackson Laboratory: Tg(Slc1a3-cre/ERT)1Nat/J) transgenic mice. We used *Nkx2.1-cre* (Xu et al., 2008) and *Cspg4-cre* (Jackson Laboratory: *B6;FVB-Tg*(*Cspg4-cre*)*1Akik/J*) (Zhu et al., 2008) transgenic mice that have been described previously. The reporter mouse *Rosa26R–Enhanced yellow fluorescent protein (EYFP*) (Srinivas et al., 2001) was used to reliably express EYFP under the control of the Rosa26 promoter upon Cre-mediated recombination.

For the induction of CreERT, Tamoxifen (20 mg/ml, Sigma, St Louis, MO) was dissolved at 37°C in 5 ml corn oil (Sigma, St Louis, MO) pre-heated at 42°C for 30 minutes. A single dose of 4 mg (250-300 µl) was administered to pregnant females by oral gavaging.

### Immunocytochemistry

Embryos were collected after caesarean section and quickly killed by decapitation. Their brains were dissected out and fixed by immersion overnight in a solution of 4% paraformaldehyde in 0.1 M phosphate buffer (pH 7.4) at 4°C. Brains were cryoprotected in a solution of 30% sucrose in 0.1 M phosphate buffer (pH 7.4), frozen and cut in 50 µm-thick coronal sections for immunostaining.

Mouse monoclonal antibodies were: BrdU (Monosan, Am. Uden, Netherlands) and GFAP (Chemicon). Rabbit polyclonal antibodies were: cleaved-caspase 3 (Chemicon), GFAP (DAKO, Carpinteria, CA); GFP (Molecular Probes, Eugene, OR); NG2 (Chemicon); Nkx2.1 (Biopat, Caserta, Italy); Olig2 (Millipore); Anti-ß galactosidase (or LacZ) (Rockland); and RFP (Labforce MBL). Guinea pig antibody was: GLAST (Chemicon, Temecula, CA). Chicken antibody was: GFP (Aves). Goat antibody was: Anti-ß galactosidase (or LacZ) (Biogenesis).

a. Fluorescence immunostaining was performed as follows: non-specific binding was blocked with 2% normal horse serum in PBS 1X solution with 0.3% Triton X-100 for preincubation and incubations. The primary antibodies were detected with Cy3-conjugated (Jackson ImmunoResearch laboratories, West Grove, PA) and Alexa488-, Alexa594- or Alexa647-conjugated antibodies (Molecular Probes, Eugene, OR). Sections were counterstained with Hoechst 33258 (Molecular Probes), mounted on glass slides and covered in Mowiol 4-88 (Calbiochem, Bad Soden, Germany).
b. DAB immunostaining was performed as follows: Endogenous peroxidase reaction was quenched with 0.5% hydrogen peroxide in methanol, and non-specific binding was blocked by adding 2% normal horse serum in Tris-buffered solutions containing 0.3% Triton X-100 for preincubation and incubations. The primary antibodies were detected with biotinylated secondary antibodies (Jackson ImmunoResearch, West Grove, PA) and the Vector-Elite ABC kit (Vector Laboratories, Burlingame, CA). The slices were mounted on glass slides, dried, dehydrated, and covered with Eukitt.

### BrdU tracing studies

To label cells in the S-phase of the cell cycle at the suitable embryonic stages (E12.5, E14.5 and E16.5), the pregnant female mice were injected intraperitoneally with a solution of 8 mg/ml of 5-bromo-2'-deoxyuridine (BrdU; Sigma, St Louis, MO) in PBS (0.15 M NaCl, 0.1 M phosphate buffer, pH = 7.4) to a final concentration of 50 mg/kg body weight. To trace the division rate of the subpallial precursors, the pregnant females were sacrificed 1-2 hours post-injection. To trace the date of genesis of the CC astrocytes, the pregnant females were sacrificed when embryos were E18.5. The BrdU was revealed by DAB or fluorescence immunostaining (as mentioned above) after a treatment with 2 M HCl for 30 min at room temperature.

### Imaging

DAB stained sections were imaged with a Zeiss Axioplan2 microscope equipped with 10×, 20× or 40× Plan neofluar objectives and coupled to a CCD camera (Axiocam MRc 1388x1040 pixels). Fluorescent-immunostained sections were imaged using confocal microscopes (Zeiss LSM 510 Meta, Leica SP5 or Zeiss LSM 710 Quasar) equipped with 10×, 20×, 40×oil Plan neofluar and 63×oil, 100×oil Plan apochromat objectives. Fluorophore excitation and scanning were done with an Argon laser 458, 488, 514 nm (blue excitation for GFP and Alexa488), with a HeNe laser 543 nm (green excitation for Alexa 594 and CY3), with a HeNe laser 633 nm (excitation for Alexa 647 and CY5) and a Diode laser 405 nm (for Hoechst-stained sections). Z-stacks of 10-15 planes were acquired for each CC coronal section in a multitrack mode avoiding crosstalk.

All 3D Z stack reconstructions and image processing were performed with Imaris 7.2.1 software. To create real 3D data sets we used the mode “Surpass”. The colocalization between two fluorochromes was calculated and visualized by creating a yellow channel. Figures were processed in Adobe Photoshop© CS4 and CS5 and schematic illustrations in Supplementary Figure 2 were produced using Adobe Illustrator© CS4.

### Quantifications

#### a) Glial cell population analysis

In 50 Im thick brain sections of *Nkx2.1*^+/+^ and *Nkx2.1*^-/-^ embryos at E18.5, the astroglial cells were labeled for GFAP and polydendroglial cells were labeled for NG2. Cells were counted in the CC, IG, MGE, MZG and POA regions from at least 4 brains per condition. The cell densities were reported per surface unit area (number of cells/mm^2^). The quantification was done using Neurolucida 9.0 and Neurolucida 9.0 Explorer© software.

In 50 Im thick brain sections of *Nkx2.1*^+/+^ and *Nkx2.1*^-/-^ embryos at E18.5, the astroglial cells that were labeled for Olig2 or both Olig2 and GLAST were counted in the CC mid from at least 2 brains per condition. Olig2 staining labeled the glial cell bodies while GLAST labeled both the cell bodies and processes. The cell densities were determined in the medial and lateral part of the CC. The cell densities were reported per volume unit (number of cells/mm^3^). The quantification was done using Imaris^®^ 7.2.1 software.

#### b) Nkx2.1^+^ and GLAST^+^ or BrdU+ cell number analyses

Pregnant female mice were injected intraperitoneally with a solution of 8 mg/ml of 5-bromo-2'-deoxyuridine in PBS to a final concentration of 50 mg/kg body weight. To trace the division rate of the subpallial precursors, the pregnant females were sacrificed 2 hours post-injection. Embryos were collected after caesarean section and quickly killed by decapitation. Their brains were dissected out and fixed by immersion overnight in a solution of 4% paraformaldehyde in 0.1 M phosphate buffer (pH 7.4) at 4°C. In 50 Cm thick brain sections of *Nkx2.1*^+/+^ and *Nkx2.1*^-/-^ embryos at E16.5, Nkx2.1^+^ cells, BrdU^+^ dividing cells and GLAST^+^ precursors or post-mitotic astroglial cells of the MGE, POA and TS were counted in the VZ, SVZ and in the parenchyma of each region, from at least 4 brains per condition. Nkx2.1 and BrdU staining labeled the cell bodies while GLAST labeled both the cell bodies and processes. The percentage of *Nkx2.1*-derived dividing precursors or post-mitotic glial cells, were determined as follows: In each sub region, and for each condition, a sample of at least four different Z-stacks was acquired at 100x magnification by using a Leica SP5 confocal microscope. The Z-stacks comprised of 10 planes that were acquired in a multitrack mode avoiding any crosstalk. Thereafter, in order to exclude the possibility of quantifying the same cells more than once, snapshots of only 3 planes (from the acquired 10 planes), were taken with Imaris^®^ 7.2.1 software (Bitplane Inc.) and analyzed.

The quantification of Nkx2.1, BrdU, GLAST and Hoechst staining was done on each snapshot separately by using Neurolucida© 9.0 and Neurolucida 9.0 Explorer© software.

#### c) Neurospheres differentiation analysis

MGE- and POA-derived neurospheres were obtained from Nkx2.1^+/+^ and Nkx2.1^-/-^ E14.5 embryos. After 7 DIV, the neurospheres were differentiated and immunostained as mentioned above. Two different brains were used for each condition and were labeled for Nkx2.1, GFAP and βIII tubulin. Cell nuclei were counterstained with Hoechst. For each condition, a total of at least 5 different Z-stacks in 5 different neurospheres were acquired at 100x magnification by using a Leica SP5 microscope. The percentage of Nkx2.1^+^/GFAP^+^ differentiated astrocytes and Nkx2.1^+^/βIII tubulin^+^ differentiated neurons were counted directly on the Z-stacks by using Imaris^®^ 7.2.1 software.

#### d) Cell death analysis

In brain sections of *Nkx2.1*^+/+^ and *Nkx2.1*^-/-^ embryos at E16.5, apoptotic cells labeled for either cleaved-caspase 3 or for TUNEL were counted in the CC, MGE, SEP, and POA from at least 2 brains per condition. 50 μm thick brain sections were used for cleaved-caspase 3 staining whereas 10 µm thick brain sections were utilized for TUNEL staining. Cell nuclei were counterstained with Hoechst. For each condition, at least 5 different Z-stacks were obtained at 100x magnification by using a Leica SP5 microscope. The number of apoptotic nuclei were counted and reported as an absolute number per section (the surface area of one section was 24119.332 µm^2^). The quantification was done using Neurolucida 9.0 and Neurolucida 9.0 Explorer© software.

### Neurosphere generation and microscopical analysis

The protocol has been adapted from Arsenijevic *et al*., 2001.

#### a) Primary culture and sphere passaging

The brains of embryos at developmental stage E14.5 were collected as described above. They were carefully removed from the skull into ice-cold sterile dissecting medium (MEM 1X) complemented with Glucose 1M (5ml/100ml). Thereafter, the brains were embedded in low melting point Agarose 3% (LMP-Agar, Gibco) at 37°C, and cut into 250 2m thick slices using a vibratome (Leica© VT 1000 S). The sections were collected in the ice-cold dissecting medium. The areas of interest (MGE, POA and SEP) were dissected out using two tungsten needles under a stereomicroscope (Leica© MZ16F). The dissected pieces of tissue were then collected into 1ml ice-cold sterile Hormone Mix Medium (MHM 1X) supplemented with Penicillin (50 U/ml) and Streptomycin (50 U/ml) (GIBCO). The Hormone Mix Medium is a growing medium containing DMEM and F-12 nutrient (1:1), glucose (0.6%), glutamine (2 mM), sodium bicarbonate (3 mM), HEPES buffer (5 mM), transferrin (100 mg/ml), insulin (25 /g/ml), progesterone (20 nM), putrescine (60 /M), selenium chloride (30 nM) (Avery et al.). Brain tissue pieces were mechanically dissociated under sterile conditions with a fire-polished pipette in the Hormone Mix Medium. The pipette was rinsed before the dissociation of each new region.

The dissociated cells were then grown in Hormone Mix Medium complemented with Pen/Strep and EGF in 6-well dishes (Nunclon Surface, NUNC Brand Products, Nalge Nunc International) at a concentration of around 10^4^-10^5^ cells per 1 ml and 4 ml per dish. After 6-7 days *in vitro* (DIV) at a temperature of 37°C in a 5% CO _2_atmosphere, the sphere cultures were expanded. Primary spheres were dissociated mechanically and cells were plated at the density of 2x10^6^ cells for 40 ml in a flask (Nunclon Surface, NUNC Brand Products, Nalge Nunc International). Sphere passages were done every 7 DIV, by spheres dissociation and transfer of 2x10^6^ cells to a new 40 ml flask.

#### b) Differentiation of spheres

After 7 DIV, the neurospheres of optimum size were chosen under a steremicroscope (Nikon©) to be transferred individually and plated onto poly-L-ornithine coated coverslips in 24-well plates (Nunclon Surface, NUNC Brand Products, Nalge Nunc International). Each coverslip contained about ten spheres and 1 ml of Hormone Mix Medium supplemented with Pen/Strep and 2% fetal bovine serum (FBS).

#### c) Immunofluorescence on differentiated Neurospheres

After 7 DIV, the neurospheres were fixed in 4% PFA for 20 minutes and permeabilized with 0.3% triton/PBS1X for 3 minutes. Coverslips were incubated with primary antibodies diluted in PBS containing 10% NHS for 2 hours at room temperature, followed by secondary fluorescent antibodies for 45 minutes at 37° and Hoechst staining for 5 minutes.

### Chromatin Immunoprecipitation

Chromatin immunoprecipitation was conducted on E16.5 brain samples according to the instructions provided by the manufacturer (Upstate, 17-295), using 2µg of a mouse anti-Nkx2.1 monoclonal antibody (MS699-P, Lab Vision). For crosslinking, 1% PFA was used. For sonication, six bursts of 45 seconds ON (30% power) and 30 second OFF were given, and samples were kept on ice during the whole sonication process. Mouse Genome Assembly data mm9 was used to map sites.

A 391 bp PCR fragment of the <u>*Lhx6* promoter</u> that includes a Nkx2.1 binding sequence at position –240 bp relative to the putative transcriptional start site was identified using primers 5’-tttgtaccgagagtaggagaagg and 5’-gtcctaactttgtagtgggcattt.

A 206 bp PCR fragment of the <u>*GFAP* promoter</u> that includes a putative Nkx2.1 binding sequence (ctcaagt) at position –838 bp relative to the putative transcriptional start site was found to be a positive binding target and was identified using primers 5’- tggataagaggccacagagg and 5’- cctctcccctgaatctctcc.

Primers against two fragments of the *Neurogenin2* promoter region, comprising of the core Nkx2.1 binding consensus sequence (tcaag), were made. 1) Primers 5’-cgggattctgactctcactaattc and 5’-aatggttctaaagctcctgttgg were designed to amplify a 410 bp PCR fragment with the core consensus Nkx2.1 binding sequence at position –668 bp relative to the putative transcriptional start site. 2) Primers 5’- cgggattctgactctcactaattc and 5’-aatggttctaaagctcctgttgg were designed to amplify another 352 bp PCR fragment with the core consensus Nkx2.1 binding sequence at position –4073 bp relative to the putative transcriptional start site.

### Transfection of HEK293 cells

A suspension of HEK-293 cells adapted to serum-free growth medium was plated at 1 x 10^6^ cells in 4 ml media in a 60mm plate. For formation of the transfection complexes, 3:1 ratio of FuGENE^®^ HD Transfection Reagent (µl): plasmid DNA (µg) was prepared and used for transfection. The study was performed by co-transfecting an expression plasmid for constitutive over-expression of Nkx2.1 *(pCAG-Nkx2.1-IRES-Tomato)* or a control plasmid *(pCAG-IRES-Tomato)* with the *pDRIVE-mGFAP* plasmid containing the GFAP promoter region in front of the LacZ reporter gene. Transfection complexes were formed by mixing 2 µg of each of the two plasmids with 12 µl of Fugene transfection reagent and 188 µl of Optimem reduced serum media. The mix was incubated at room temperature for 20 minutes and thereafter, added to the cell plates. The cell plates were kept in the 37 °C incubator and gene expression analysis was done after 24-48 hours of transfection. Fluorescence immunostaining was done to visualize the presence and level of *LacZ* expression. Tomato signal was visible by direct fluorescence, however, for a clearer visualization of Tomato signal, an anti-RFP immunostaining was done. The method to do fluorescence immunostaining has been described above.

### Statistical analysis

The results from all quantifications were analyzed with the aid of Statview software (SAS Institute Inc.). For all analysis, values from at least three independent experiments were first tested for normality and the variance of independent populations were tested for equality. Values that followed a normal distribution were compared using Student's *t*-test. Values that did not follow a normal distribution were compared using Mann-Whitney non-parametric test.

### Atlas and nomenclature

The neuroanatomical nomenclature is based on the “Atlas of the prenatal mouse brain” (Schambra et al., 1991).

## Abbreviations list

AEP: Anterior entopeduncular area
BrdU: 5-bromo-2’-deoxyuridine
CC: Corpus callosum
CCi: Cingulate cortex
CI: Cingulate bundle
Cre-ERT^tm^: Tamoxifen inducible Cre recombinase fused to the ligand binding domain of the estrogen receptor
Cspg4: Chondroitin sulfate proteoglycan 4 (also known as NG2)
DIV: Day in vitro
E: Embryonic day
EGFP: enhanced green fluorescent protein
EYFP: enhanced yellow fluorescent protein
GABAergic: γ-aminobutyric acidergic
Gad1: Glutamate decarboxylase 1 (also known as GAD67)
*Gad1-EGFP*: Gad1-EGFP knock-in mouse
GFAP: Glial fibrillary acidic protein
GLAST: Glutamate-aspartate transporter
GP: Globus pallidus
HIC: Hippocampal commissure
IG: Indusium griseum
IRES: Internal ribosome entry site
IZ: Intermediate zone
KO: knockout
LGE: Lateral ganglionic eminence
LH: Lateral hypothalmus
Lhx6: LIM homeodomain (LIM-hd) gene
LPOA: lateral preoptic area
LV: Lateral ventricle
MGE: Medial ganglionic eminence
MZ: Marginal zone
MZG: Midline zipper glia
NG2: Neuron-glial antigen 2
Ngn2: Neurogenin2
*Nkx2.1*: NK2 homeobox 1
*Nkx2.1-Cre*: Mouse with the Cre recombinase under control of the *Nkx2.1* promoter
Olig2: Oligodendrocyte transcription factor
pCAG: promoter constructed from following sequences: (**C**) cytomegalovirus early enhancer element, (**A**) promoter, the first exon and intron of chicken beta-actin gene, (**G**) the splice acceptor of the rabbit beta-globin gene
POA: Preoptic area
RMS: Rostral migratory stream
*Rosa-EYFP*: Rosa26-lox-STOP-lox-EYFP reporter mouse
S100ß: Small EF-hand calcium and zinc binding protein
SEP: Septum
ST: Striatum
SVZ: Subventricular zone
TS: Triangular septal nucleus
TUNEL: Terminal deoxynucleotidyl transferase dUTP nick end labeling
VZ: Ventricular zone
WT: wild-type

## Acknowledgements

We are particularly grateful to Christiane Devenoges for technical assistance. We would like to thank F. Thevenaz and Alain Gnecchi for mouse care, plugs and genotyping. We thank Jean-Yves Chatton from the Cellular Imaging Facility (CIF, University of Lausanne, Switzerland) for imaging assistance. Shilpi Minocha was supported by a postdoctoral fellowship of the Fondation Pierre Mercier pour la science. The work in the laboratory of C. Lebrand was supported by funds from Swiss National Foundation Grant # 31003A-122550.

---

**Figure 1-figure supplement 1.**
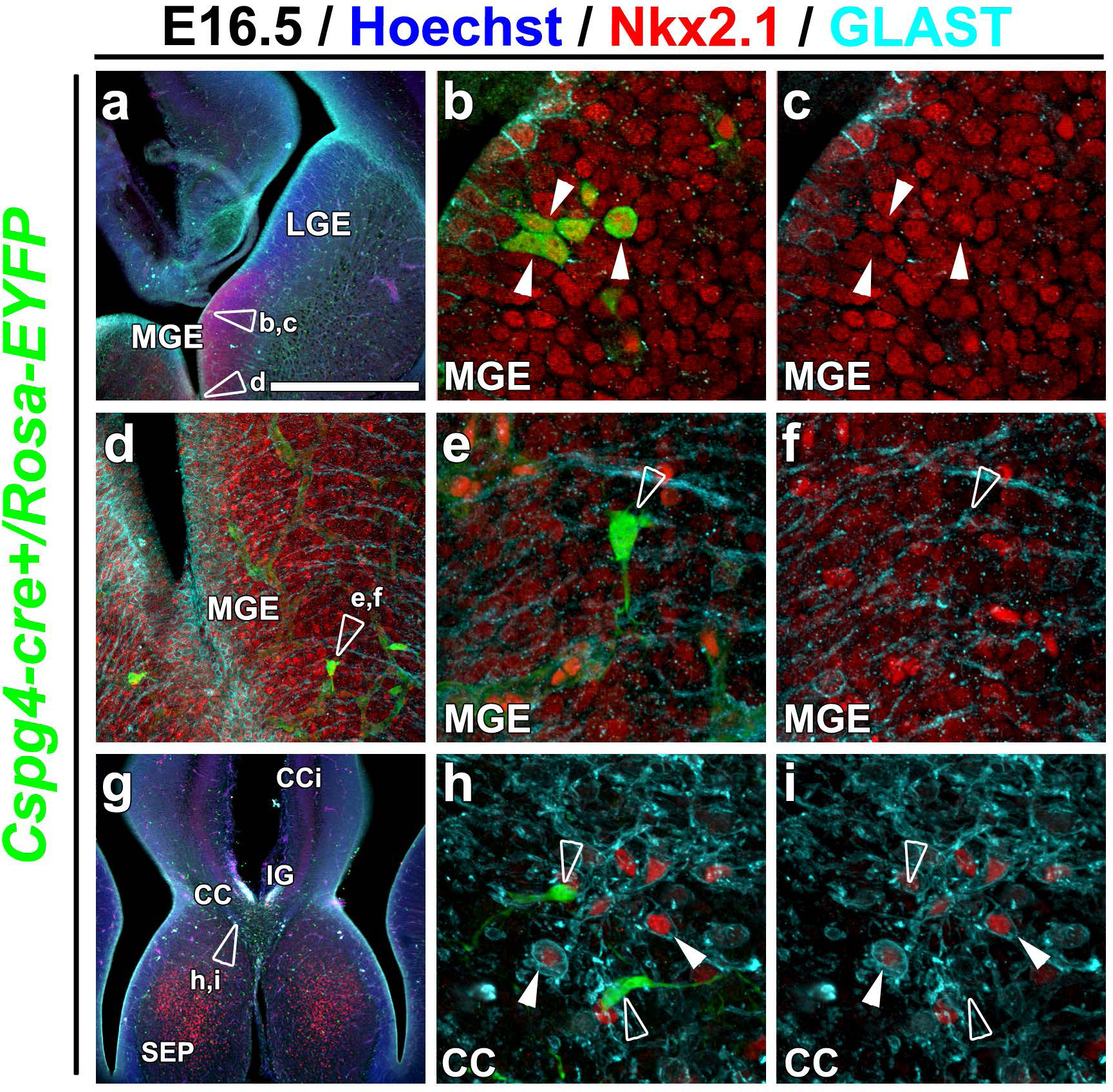
Fate-mapping study of *Nkx2.1*-regulated NG2^+^ polydendrocytes using the *Cspg4-Cre^+^/Rosa-EYFP* reporter mice. **(a-i)** Triple immunohistochemistry for the EYFP, Nkx2.1 and GLAST on coronal sections from *Cspg4-Cre^+^/Rosa-EYFP* mice (n=3) at E16.5. Cell nuclei were counterstained in blue with Hoechst **(a** and **g)**. **b, c, e, f, h** and **i** are higher power views of the regions shown in **a, d** and **g**, respectively. At E16.5, NG2^+^ (or Cspg4^+^) polydendrocytes visualized by the EYFP signal were found to originate from Nkx2.1^+^ subpallial sites such as the MGE **(a-c** and **d-f)**. The colocalization between Nkx2.1 (in red) and the EYFP signal (in green) is observed in few cells in the SVZ of the MGE (solid arrowheads in **b** and **c**) but as soon as the NG2^+^ cells start to differentiate and migrate, Nkx2.1 is down-regulated and is no anymore more visible (open arrowheads in **e-f** and **h-i**). By contrast, Nkx2.1 is still expressed in GLAST^+^ astroglial cells within the CC midline (solid arrowheads in **h-i**). **(CC)** corpus callosum; **(CCi)** cingulate cortex; **(CI)** cingulate bundle; **(IG)** induseum griseum; **(MGE)** medial ganglionic eminence; **(SEP)** septum. Bar = 675 µm in **a** and **g**; 160 µm in **d**, 40 µm in **b, c, e, f, h** and **i**.

**Figure 1-figure supplement 2.**
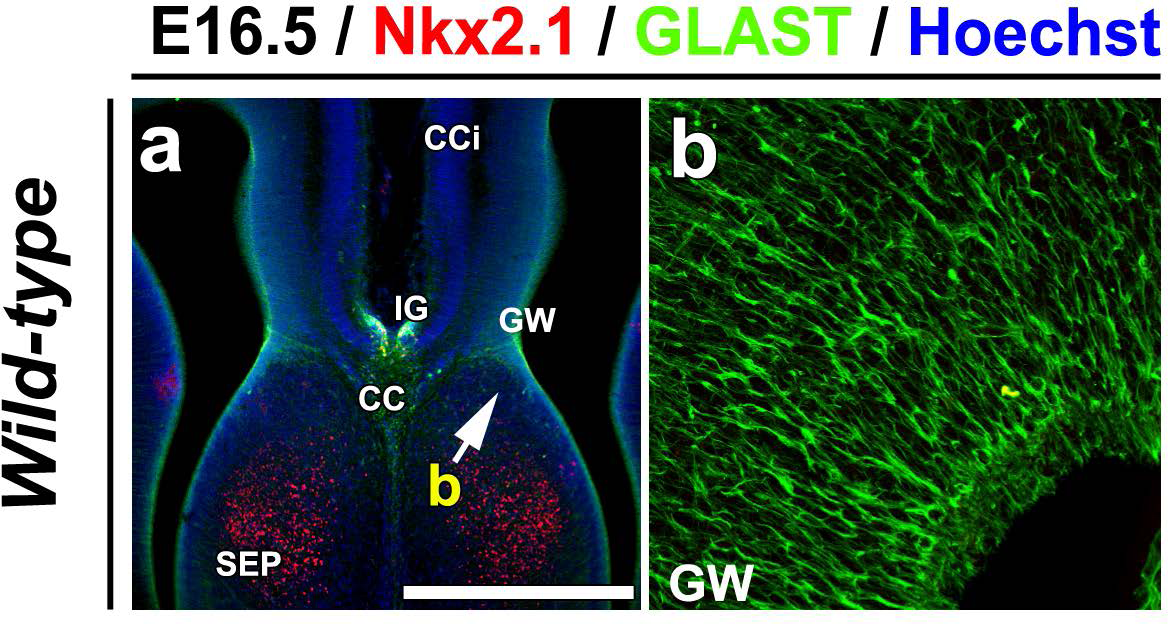
Radial glial cells within glial wedge region are Nkx2.1-negative. **(a-b)** Double immunostaining for the Nkx2.1 and GLAST on coronal sections from WT mice (n=3) at E16.5. Cell nuclei were counterstained in blue with Hoechst **(a)**. **b** is higher power view of the regions shown in **a**. The GLAST^+^ radial glial cells within glial wedge **(GW)** do not express Nkx2.1. **(CC)** corpus callosum; **(CCi)** cingulate cortex; **(IG)** induseum griseum; **(SEP)** septum. Bar = 675 µm in **a** and 40 µm in **b.**

**Figure 1-figure supplement 3.**
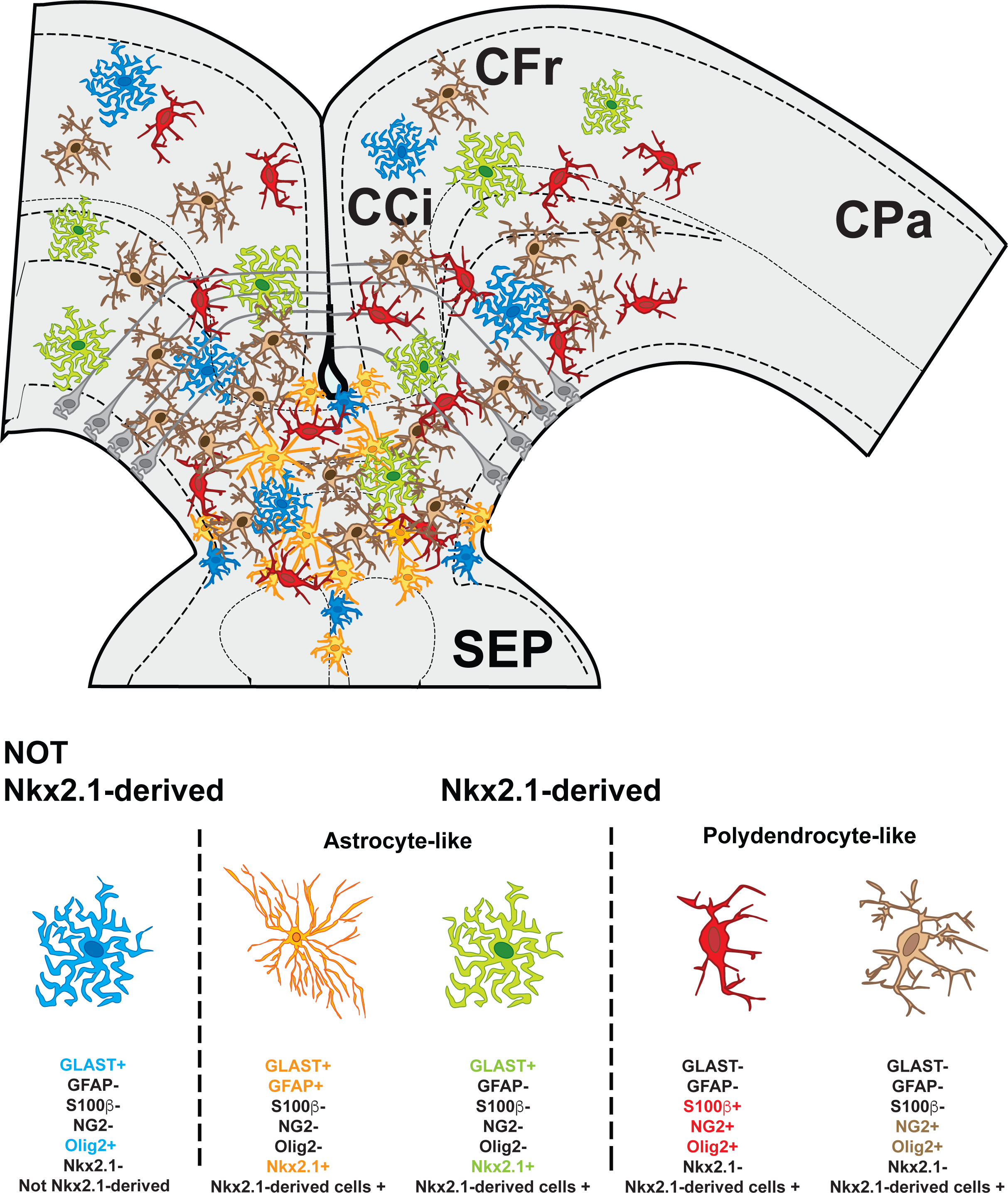
Four subtypes of *Nkx2.1*-derived glial cells. The schema represents a coronal view of the CC at E18.5, and summarizes the different types of *Nkx2.1*-derived glial populations visualized in our experiments. The CC forms a complex environment composed of one non-*Nkx2.1*-derived glial cell subtype and four different subtypes of *Nkx2.1*-derived glial cells. Three types of astrocyte-like cell populations are shown: in orange, the GLAST^+^/GFAP^+^/Olig2-/Nkx2.1^+^ cells, in green, the GLAST^+^/GFAP^-^/Olig2^-^/Nkx2.1^+^ cells, and in blue, the GLAST^+^/Olig2^+^/Nkx2.1^-^ cells. Two types of polydendrocyte-like cells are shown; in red, the GLAST^-^/S100β^+^/NG2^+^/Olig2^+^/Nkx2.1^-^ cells, and in brown, the GLAST-/NG2^+^/Olig2^+^/Nkx2.1^-^ cells. Under each glial cell-type category, the expression profile of the different glial markers, employed to identify and characterize the glial cells, used in combination with the Nkx2.1 antibody, is presented. The (**+**) sign indicates that the glial cell type was positively labelled by the listed marker, whereas the (–) sign indicates that the glial cell type was not labelled by the listed marker. **(CCi)** cingulate cortex, **(CFr)** frontal cortex, **(CPa)** parietal cortex, **(SEP)** septum.

**Figure 4-figure supplement 1.**
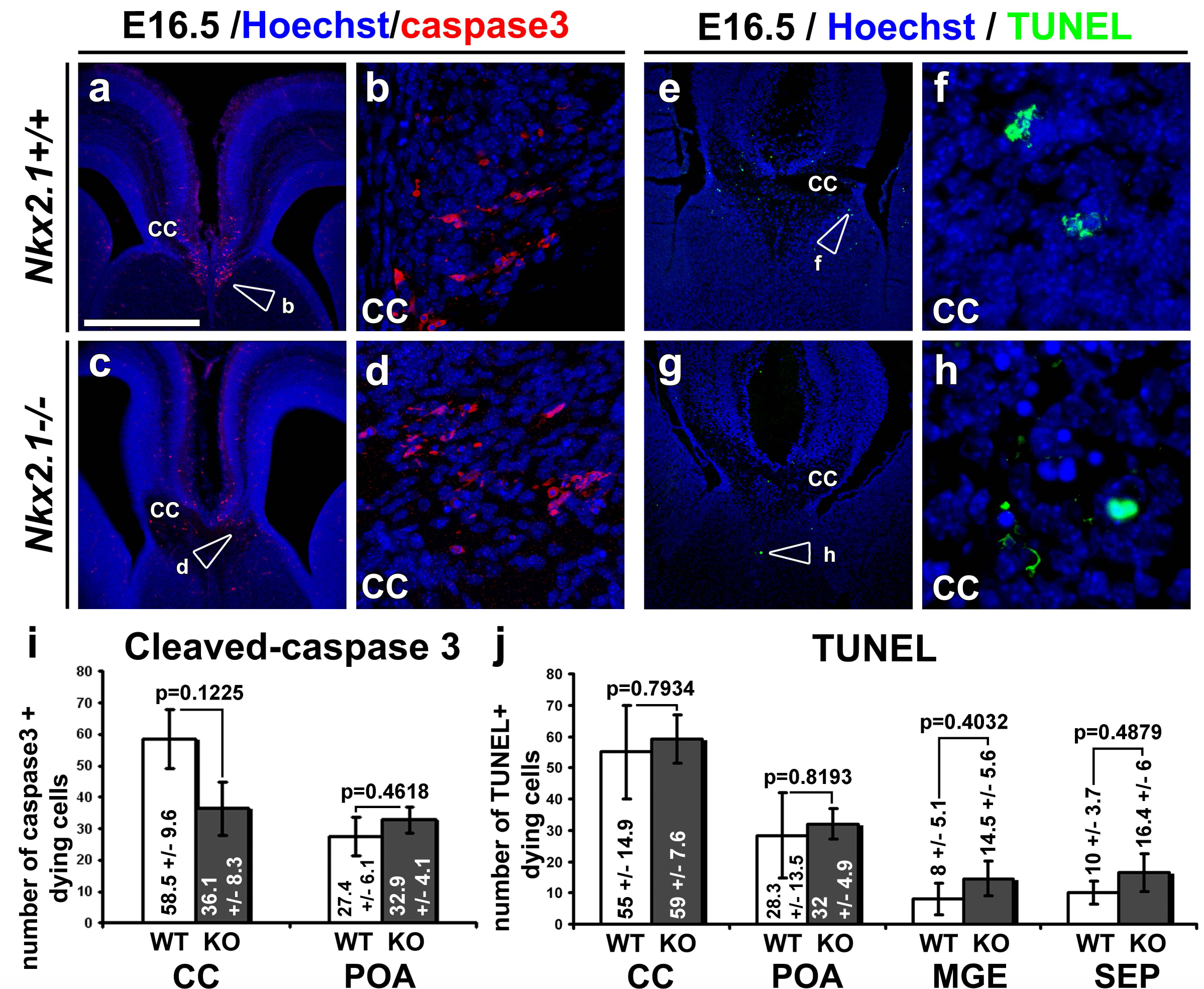
*Nkx2.1^-/-^* mice brains do not show any increase in cell death at E16.5. **(a-d)** Single immunohistochemical staining for the cleaved-caspase 3 (n=4 for CC region and n= 5 for POA region in WT mice; n=6 for CC region and n=10 for POA region in *Nkx2.1^-/-^* mice) and **(e-h)** TUNEL staining (n=16 for CC in WT mice, n=22 for CC in *Nkx2.1^-/-^* mice; n=6 for POA in WT mice, n=5 for POA in *Nkx2.1^-/-^* mice; n=10 for MGE in WT mice, n= 11 for MGE in *Nkx2.1^-/-^* mice; n=7 for SEP in WT mice, n=14 for SEP in *Nkx2.1^-/-^* mice) on CC coronal sections from wild-type **(a-b** and **e-f)** and *Nkx2.1^-/-^* mice **(c-d** and **g-h)** at E16.5. Cell nuclei were counterstained in blue with Hoechst. **b**, **d**, **f** and **h** are higher magnified views of the CC region seen in **a**, **c**, **e** and **g**, respectively. **(i** and **j)** Bars (mean ± SEM from a sample of n=4-16 sections in the wild-type and n=5-22 sections in *Nkx2.1^-/-^* mice depending on the region studied) represent the number of dying cells labelled by the cleaved-caspase 3 or by the TUNEL staining and displaying pyknotic nuclei per section (surface area/section=24119.332 mm^2^), in the CC, POA, MGE and SEP of *Nkx2.1^-/-^* (KO) compared to wild-type (WT) mice. No significant differences were observed in the number of dying cells in *Nkx2.1^-/-^* mice brains compared to the wildtype. p-value= 0.1225 for CC and 0.4618 for POA with cleaved caspase 3 staining. p-value= 0.7934 for CC, 0.8193 for POA, 0.4032 for MGE, and 0.4879 for SEP with TUNEL staining. **(CC)** corpus callosum; **(MGE)** medial ganglionic eminence; **(POA)** preoptic area; **(SEP)** septum. Bar = 675 µm in **a**, **c**, **e** and **g**; 60 µm in **b** and **d**; 40 µm in **f** and **h**.

